# All-atom simulations elucidate the molecular mechanism underlying RNA-membrane interactions

**DOI:** 10.1101/2024.11.01.618995

**Authors:** Salvatore Di Marco, Jana Aupič, Giovanni Bussi, Alessandra Magistrato

## Abstract

RNA-membrane interactions are starting to emerge as an important organizing force in both natural and synthetic biological systems. Notably, RNA molecules were recently discovered to be present on the extracellular surface of living cells, where they mediate intercellular signalling. Furthermore, RNA-membrane interactions influence the efficacy of lipid-based RNA delivery systems. However, the molecular terms driving RNA localisation at the membrane remain poorly understood. In this work, we investigate how RNA-phospholipid membrane interactions occur, by means of all-atom simulations. We find that among the four RNA nucleobases guanine exhibits the strongest interaction with the membrane due to extensive hydrogen bond formation. Additionally, we show that intra-RNA base pairing present in organised RNA structures significantly hinders RNA binding to the membrane. Elucidating the molecular details of RNA-membrane association will importantly contribute to improving the design of RNA-based drugs as well as lipid-based RNA delivery systems and to parsing out RNA transport and localisation mechanisms.

## Introduction

RNA is a fundamental and versatile molecule in biology, capable of storing genetic information, catalyzing chemical reactions and regulating diverse cellular processes such as DNA transcription and protein synthesis. It recently came into view that RNA molecules also participate in cell-to-cell communication. For example, RNA molecules can be found tethered on the surface of cell membranes [1, 2]. These RNAs, some decorated with glycans, play key roles in intercellular signalling, and were shown to influence immune cell differentiation and breast cancer transformation [3]. Alternatively, RNA molecules like mRNA and miRNA, can be encapsulated into extracellular vesicles (EVs) via different pathways, some of them lipid-mediated, and subsequently transmitted to other cells [4, 5]. While the molecular mechanisms governing RNA transport, its anchoring to the membrane surface and encapsulation into EVs are still obscure, these studies indicate that RNA-membrane association is pivotal for RNA-mediated signalling.

Furthermore, RNA-membrane interactions are also relevant for synthetic biology. The most prominent example is the development of new RNA-based therapeutics, since RNA molecules must be packaged into lipid nanoparticles (LNPs), to shield them from external factors and to achieve RNA delivery across cellular membranes [6]. LNPs contain a mixture of a phospholipids, ionisable lipids, PEG-ylated lipids and cholesterol [6]. Their composition importantly affects LNP formulation stability, RNA intracellular delivery as well as the immunogenic response and must thus be carefully optimised. At present this is accomplished empirically by laborious screening of large numbers of different LNPs, as there are no established design principles primarily due to limited comprehension of the molecular processes that underlie the packing of RNA molecules into LNPs and their cytosolic delivery through endocytosis [6–8].

Lastly, RNA-membrane interactions may also have played a role in the origin of life [9, 10]. According to the RNA World theory cellular organisms emerged from compartmentalised self-replicating RNA molecules. Indeed, RNA-membrane interactions were shown to modulate both membrane permeability [11] and activity of catalytic RNAs [12], giving further credence to the RNA World hypothesis.

Despite their emerging relevance for both natural and synthetic systems, RNA-lipid bilayer interactions have been hitherto largely overlooked. Early experimental studies demonstrated that the binding of RNA molecules to membrane systems is influenced by several factors. Divalent cations were shown to significantly enhance RNA-membrane interactions [13]. Conversely, increasing ionic strength by adding monovalent salts reduced RNA-lipid binding [14, 15]. Furthermore, the strength of the interaction was shown to depend also on RNA sequence, with guanine (G) rich RNAs exhibiting tighter binding [12]. Conversely, the role of structure is less clear with some studies reporting preferential binding of ssRNA [16], while others observed a stronger interaction with dsRNA [10, 17, 18].

All-atom molecular dynamics (MD) simulations represent a powerful tool for investigating RNA-membrane interactions, since their dependence on RNA sequence and structure can be dissected on a molecular level under precisely defined solution conditions [19–21]. Coupled with enhanced sampling methods [22], that speed up the exploration of rare events, all-atom simulations also enable precise determination of energetics associated with various molecular processes, such as RNA binding to membrane.

Here, we employ metadynamics [23], an enhanced sampling method, to systematically examine the molecular principles underlying RNA interactions with a fluid phospholipid bilayer. We determine the binding free energy for RNA sequences of different length and composition, starting from a single nucleoside to a 19-mer strand with either defined or undefined structural features. We demonstrate guanosine has the highest affinity for the lipid bilayer, with hydrogen bond formation as the primary driving force for RNA-membrane association. Consequently, structured RNA molecules, with nucleobases engaged in base pairing, were observed to interact less strongly with the lipid bilayer than unstructured RNA strands of comparable length. Taken together, our simulations provide important groundwork for studying mechanisms of RNA-membrane association in living systems and may aid in the design of RNA drugs and lipid-based RNA drug delivery systems.

## Results

### Purines exhibit a higher affinity for the membrane than pyrimidines

First, we set out to characterise and compare the binding modes and affinities of different nucleosides to the phospholipid bilayer. Our model membrane was constructed from dipalmitoylphosphatidylcholine (DPPC) lipids. In all the studied systems, Amber force fields *χ*OL3 [24–26] and Lipid21 [27] were employed to describe the RNA and membrane components, respectively. To enhance the sampling of the nucleoside binding/unbinding events, we performed well-tempered metadynamics (wtMD) [28] utilizing the absolute value of the z-distance |*d_z_*| from the membrane as a collective variable (CV). Details about the definition of this quantity are presented in the Methods section. Each simulation was run for 4 *µ*s, permitting us to observe several binding and unbinding events for each nucleoside and thus reliably determine the associated free energy profiles (Fig. 1a). The free energy minimum for every nucleoside, except adenosine (A), was observed around 1.6–1.7 nm from the membrane center, corresponding to the membrane surface. Conversely, in case of A the lowest energy states were observed at *d_z_* ≈ 1.4 nm, indicating that A preferred to penetrate into the polar headgroup layer of the membrane. The free energy difference Δ*F* between unbound and bound states was around 20 kJ/mol for purines (G and A) and around 12 kJ/mol for cytosine (C) and uracil (U). In all cases, membrane binding was a barrierless process.

**Figure 1:**
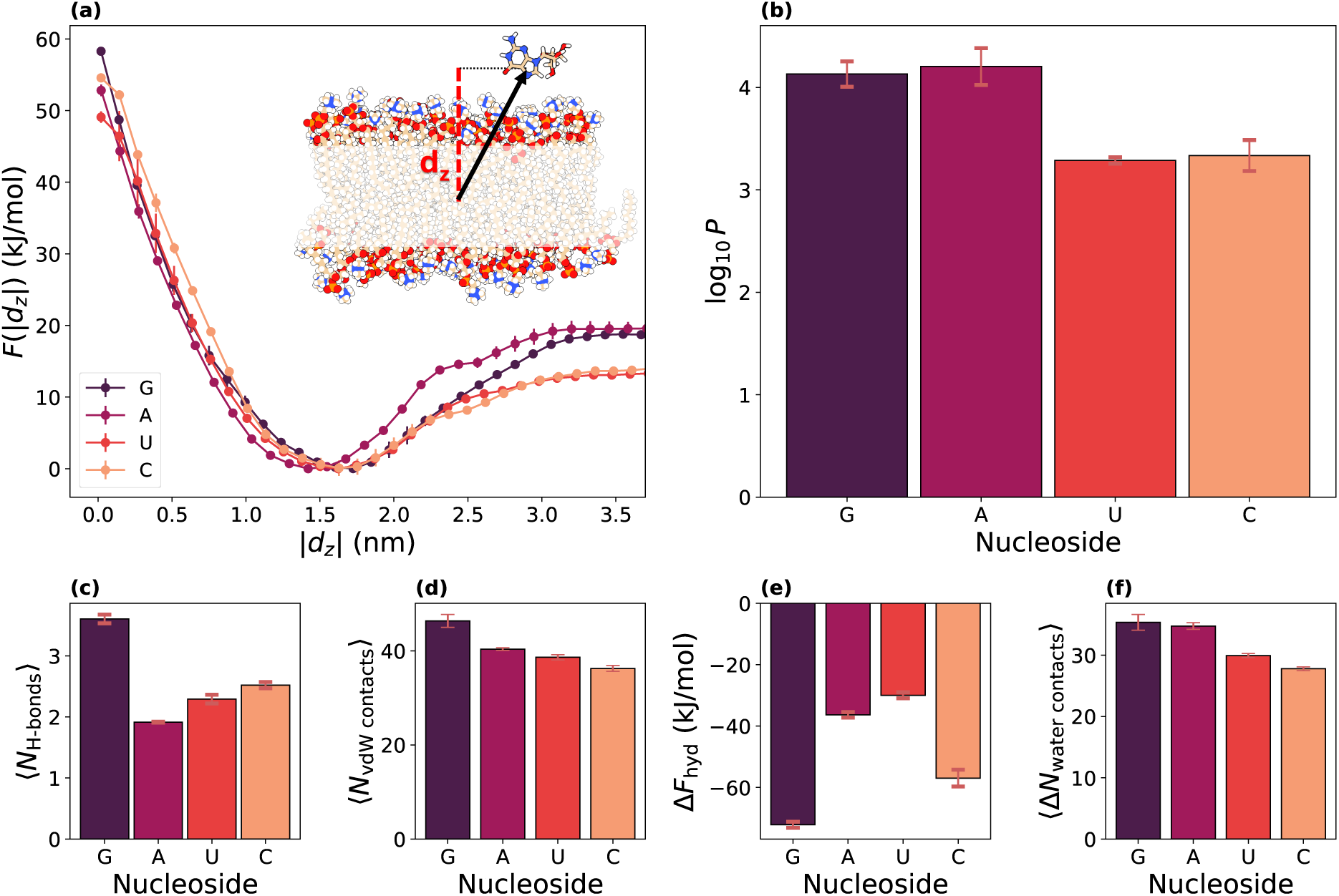
Binding affinity of RNA nucleosides to the fluid lipid bilayer. (a) Interaction free energy *F* as a function of the absolute value of the z-component of the distance between nucleoside and membrane centers (|*d_z_*|). A representation of a simulated system, highlighting the definition of the collective variable *d_z_*, is displayed as inset. (b) Base 10 logarithm of the partition coefficient (log_10_ *P*) for each nucleoside. (c) Average number of hydrogen bonds with the membrane (⟨*N*_H-bonds_⟩) for each nucleoside in the bound state. (d) Average number of vdW contacts between heavy atoms of each nucleoside and the membrane (⟨*N*_vdW contacts_⟩) in the bound state. (e) Hydration free energies (Δ*F*_hyd_) estimated via 3D-RISM. (f) Average number of vdW water contacts lost upon membrane binding (⟨Δ*N*_water contacts_⟩) for each nucleoside.

Additionally, to facilitate comparison to experimental data, we computed partition coefficients (*P* s, Fig. 1b). *P* s, related to Δ*F*, indicate how likely it is for a molecule to be in the bound versus the unbound state (see Methods). A high *P* implies that a molecule is more likely to be located in the membrane partition than in bulk solvent. As expected from the free energy profiles, purines were characterised by a higher *P* (log_10_ *P* ≈ 4) than pyrimidines (log_10_ *P* ≈ 3).

To explain the observed differences in behaviour, we analyzed the formation of hydrogen bonds and van der Waals (vdW) contacts between the nucleosides and the membrane in the bound state, and estimated the former’s hydration free energy. Since DPPC lipids can only accept hydrogen bonds, nucleosides must act as hydrogen bond donors (Fig. 1b). While all nucleosides can be involved in hydrogen bonding via their ribose moiety in equal measure (Fig. S1), the nucleobases diverge in the number of hydrogen bond donating moieties. G can contribute as much as three protons to hydrogen bonding. Instead, A and C can each donate two, originating from the amino group, while U is able of donating a single proton. Indeed, G formed a significantly higher number of hydrogen bonds (3.6 ± 0.1) than other nucleosides (<2.5). Interestingly, despite having a similar *P* as G, A was only poorly involved in hydrogen bonding. To incorporate also other forms of interactions, we then analysed vdW contacts (Fig. 1d). Surprisingly, though A had a tendency to wedge inside the membrane, G again formed the highest amount of vdW contacts. Instead, A displayed an interaction profile that was comparable to those observed for C and U, which are characterised by a 10-times lower P. Seeking to rationalise the high *P* observed for A, we computed nucleosides hydration free energies using 3D-RISM [29] (Fig. 1e). We remark that although the hydration free energy estimated with this method are in general [30] markedly overestimated, here we aim to compute the differences between the different bases. As a result, G and C had the highest hydration free energy, while A and U were significantly more hydrophobic, in accordance to previous experimental and computational works [31–33]. This suggests that A’s high affinity for the membrane layer is driven by disruption of its unfavourable interactions with water. Accordingly, analysis of water vdW contacts in the unbound and bound states demonstrated that A lost significantly more vdW contacts with water upon membrane binding than U, which is characterised by a similar hydrophobicity but lower *P* (Fig. 1f).

Lastly, we investigated the orientation of nucleobases once bound to the membrane surface. For this purpose, we calculated the orientational free energy *F* (cos(*θ*)), where *θ* is defined as the angle between the normal of the nucleobase and the membrane normal, aligned with the z-axis (Fig. S2, Fig. 2). Although orientational free energy differences were small, we could identify subtle differences. G preferred to be stacked on top of the membrane and not inserted as it can be observed from the free energy maximum at 0 (Fig. S2). Conversely, A bound perpendicularly to the surface, while C and U had less defined orientation preferences. It is important to note we did not observe symmetric behavior with respect to cos(*θ*) = 0, since nucleobases are attached to the ribose. Consequently, depending on the nucleobase side that interacts with the membrane surface, the ribose will be positioned differently, leading to free energy asymmetries with respect to cos(*θ*) sign inversion.

**Figure 2:**
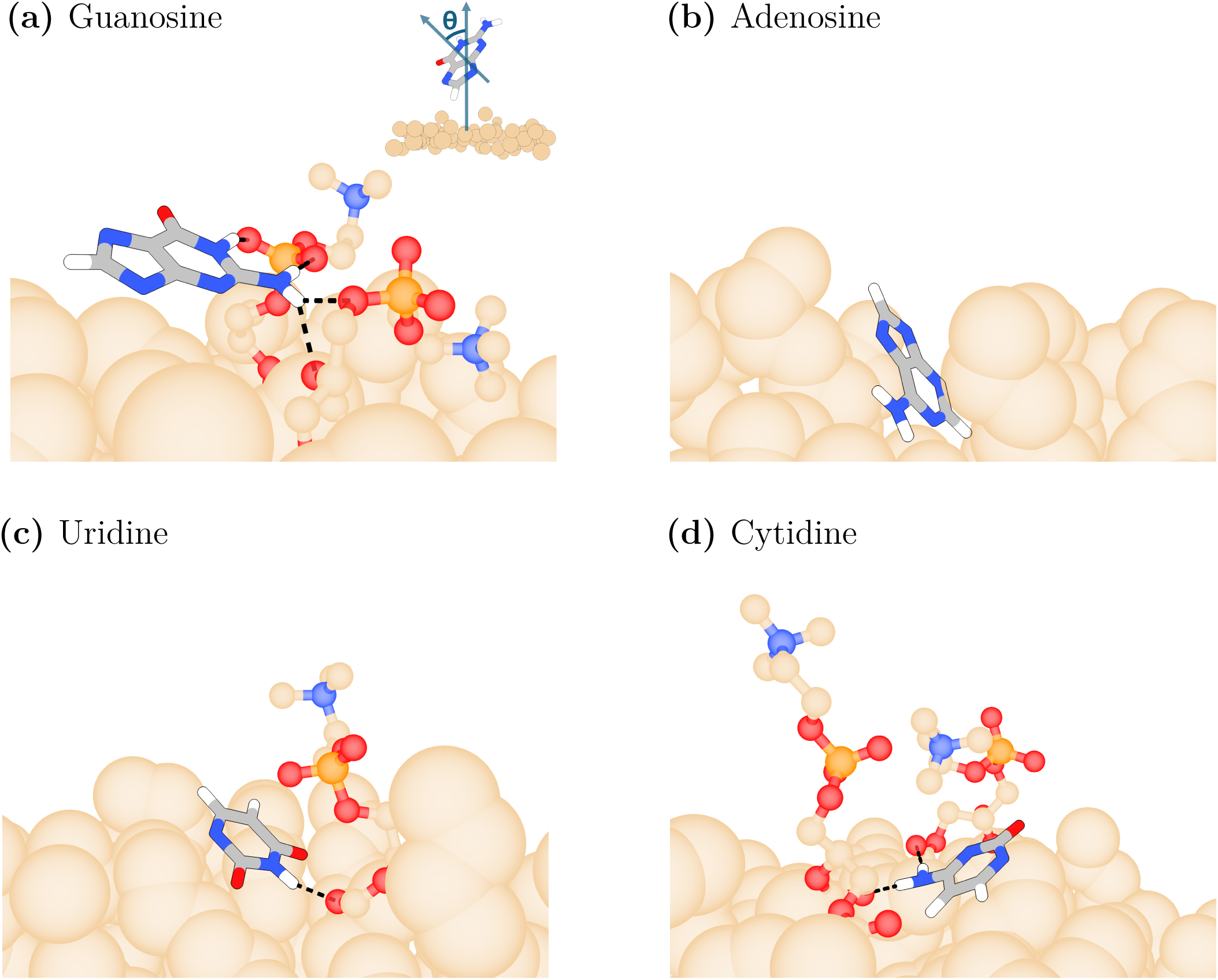
Nucleosides exhibit distinct binding modes in the membrane-bound state. Preferred orientation for each nucleoside ((a) guanosine, (b) adenosine, (c) uridine and (d) cytidine) was obtained by analyzing the free energy in the bound state as a function of the angle (*θ*) between the nucleobase and membrane normal. The definition of *θ* is depicted in the inset of panel a. Representative frames are shown. For clarity, ribose rings are not shown and only lipid head groups hydrogen bonding to nucleosides are displayed. Dashed black lines represent hydrogen bonds between nucleosides and the membrane. Adenosine orients perpendicularly to the membrane and is inserted deeper, while other nucleosides tend to adjust in a roughly parallel way.

### Guanine homopolynucleotides display the highest binding affinity

Next, we examined how membrane binding is affected by RNA sequence length by simulating homopolynucleotides composed of 2 or 3 nucleotides. In contrast to single nucleosides, these systems contained one or two negatively charged phosphate groups that could alter the observed binding modes and affinities. Consistently with the affinity rankings obtained for nucleotides, we observed that guanine dinucleotide (G2) and trinucleotide (G3) had a higher partition coefficient than other sequences with the same number of nucleotides (Figs. 3a, S3, S4) G is also the only nucleotide where *P* systematically increased with sequence length (Fig. 3a). This is due to G consistently forming more hydrogen bonds and vdW contacts with the membrane surface as the sequence is made longer, while other nucleobases showed only a moderate increase, if any (Figs. 3b, 3c). Interestingly, in all cases we observed a decrease in the average total number of vdW contacts per nucleotide and a significant reduction in hydrogen bonds formed via the ribose sugars (Figs. S5, S6). This is likely a result of the reduced conformational freedom due to the linking of nucleotides into a single chain and the presence of phosphate groups that represent a sterical hindrance, both of which prevent the nucleobases to assume their preferred binding poses. Indeed, we observed that phosphate groups prefer to orient away from the membrane, maintaining contact with water molecules and DPPC lipid choline moiety (Figs. S7, S8, S9). While the latter interaction represent a favourable contribution to RNA-membrane association it is not sufficiently strong to offset the loss of hydrogen bonding between the nucleobases and the membrane. Additionally, in case of the polyA sequences the reduced hydrophobicity due to the increased backbone charge further lowers partitioning of A residues into the lipid bilayer.

**Figure 3:**
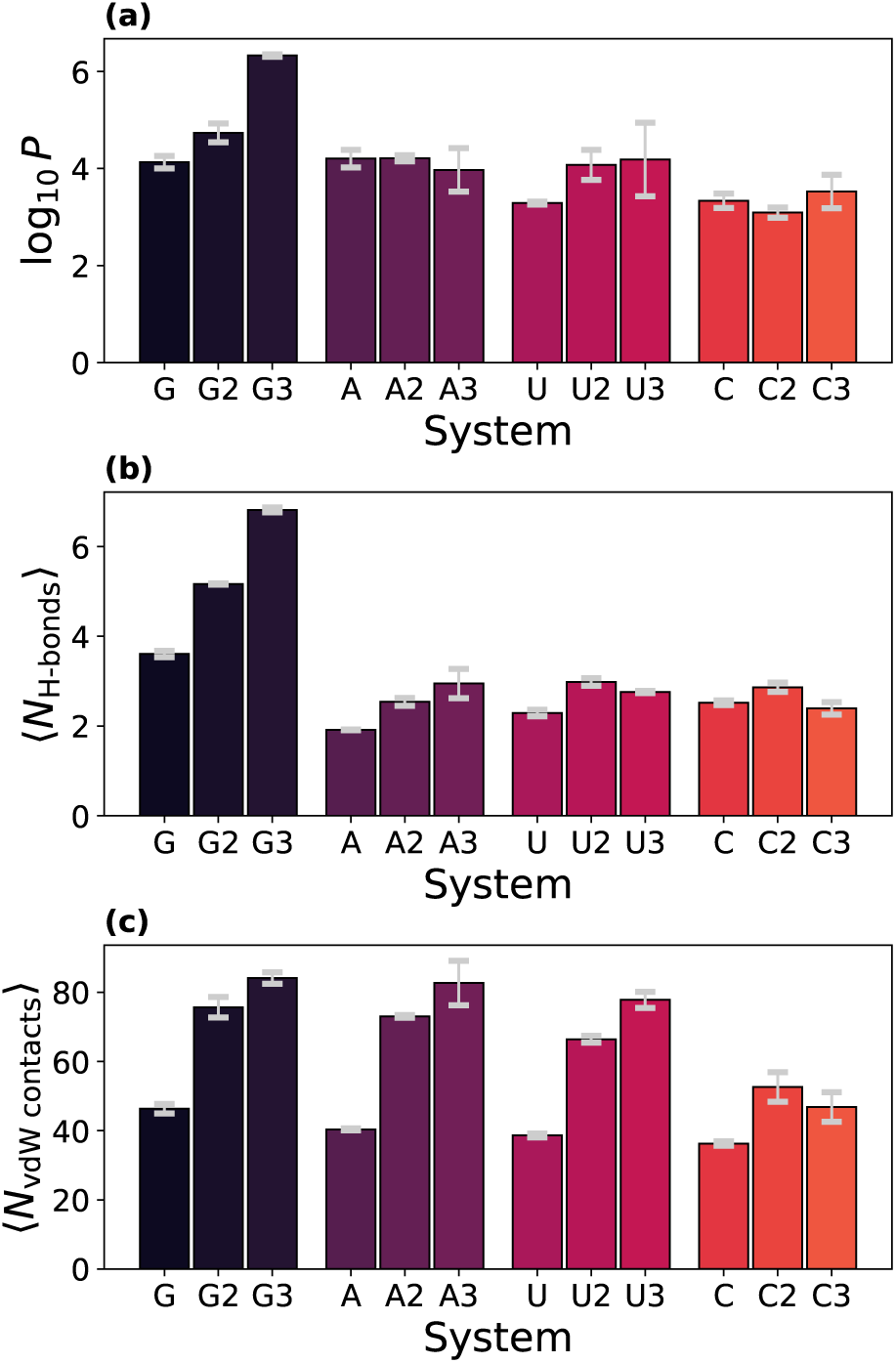
Affinity of polyguanine sequences for the lipid bilayer increases with sequence length. (a) Logarithm of the partition coefficients (log_10_ *P*) for the different studied nucleoside and polynucleotide sequences (N; nucleoside, N2; dinucleotide, N3; trinucleotide). (b) Average number of hydrogen bonds made by each nucleoside or polynucleotide with the membrane (⟨*N*_H-bonds_⟩) in their bound state. (c) Average number of van der Waals contacts with the membrane (⟨*N*_vdW contacts_⟩) made by each nucleoside or polynucleotide in their bound state.

**Figure 4:**
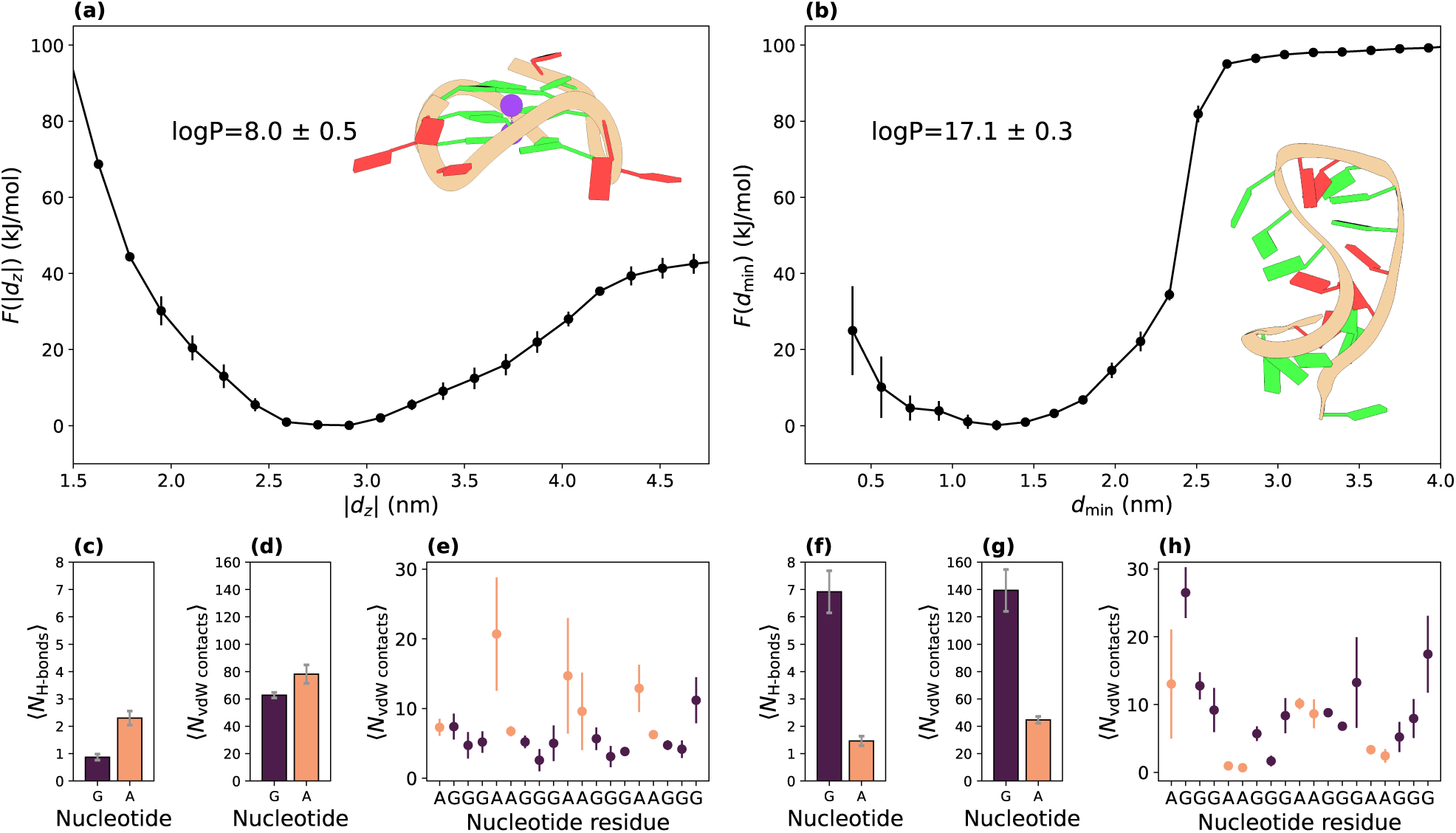
Structure inhibits interaction. (a) Free energy (*F*) as a function of the absolute value of the z-distance between the RNA G-quadruplex core and the center of the membrane (|*d_z_*|). (b) *F* as a function of the minimum distance between the unfolded RNA strand and the membrane center (*d*_min_). (c) Average number of hydrogen bonds (⟨*N*_H-bonds_⟩) formed between the membrane and the G-quadruplex in the bound state for each nucleotide type. (d) Average number of van der Waals contacts (⟨*N*_vdW contacts_⟩) between the membrane and the G-quadruplex in the bound state for each nucleotide type. (e) ⟨*N*_vdW contacts_⟩ with the membrane for each nucleotide residue in the G-quadruplex. (f) ⟨*N*_H-bonds_⟩ formed between the membrane and the unfolded RNA strand in the bound state for each nucleotide type. (g) ⟨*N*_vdW contacts_⟩ between the membrane and the unfolded RNA strand in the bound state for each nucleotide type. (h) ⟨*N*_vdW contacts_⟩ with the membrane for each nucleotide residue in the unfolded RNA strand.

Analysis of binding modes revealed that in G-, C- and U-based sequences, nucleobases preferred to orient parallel to the membrane surface (Figs. S10, S11). Conversely, for A-based sequences, we observed more heterogeneous binding modes, where, in the majority of cases, at least one nucleobase oriented perpendicular to the membrane (Figs. S10, S11). Additionally, for all nucleotides, no significant differences in conformational and base stacking behaviour were observed between bound and unbound states.

### RNA folding hinders interaction with the membrane

Finally, we attempted to parse the effect of RNA folding on RNA-membrane association. To this end, we simulated a G-rich 19-mer RNA sequence both in an unfolded and a folded, G-quadruplex conformation. The G-quadruplex was composed of a guanine core held in place by two K^+^ ions, while its loops contained purely A residues. This RNA sequence was chosen since its interaction with the membrane was assessed experimentally in previous studies [12]. Due to the more complicated nature of the simulated systems, we used two CVs per system. For the G-quadruplex, we employed as CVs the G-quadruplex orientation with respect to the membrane normal and the z-distance between the G-quadruplex core and the membrane center (*d_z_*). For the unfolded RNA strand, we instead used the maximum and minimum distances between the RNA strand and the membrane surface (*d*_max_ and *d*_min_, respectively). Details are explained in the Methods section.

The unfolded RNA 19-mer displayed a significantly higher affinity for the lipid bilayer than the folded G-quadruplex, exhibiting a two-fold decrease in Δ*F* (Figs. 4a, 4b). Indeed, the unfolded RNA strand formed more than 2.5 times the average number of hydrogen bonds and approximately 30% more vdW contacts than the G-quadruplex, explaining the large difference in computed *P* s and Δ*F* s (Figs. 4(c–h), S12). The lower engagement of the G-quadruplex in membrane binding was expected, since G residues making up the G-quadruplex core are not available for interaction with the membrane (Figs. 4(c–h), S12). Indeed, looking at the number of vdW contacts made by each nucleotide in the sequence, we observed that in the G-quadruplex state only the loop regions composed of As interacted with the membrane, while in the unfolded conformation, the G residues drove membrane association (Figs. 4e, 4h).

Next, we identified the preferred binding modes of the two systems. The G-quadruplex structure bound by stacking on top of the membrane, maximizing contacts between the membrane and A residues located in the loop regions (Figs. S13, S14). Conversely, the free energy minimum for the unfolded RNA strand corresponds to conformations where the RNA is fully adsorbed on the membrane with most nucleotides making contact with the membrane (Figs. S15,S16).

## Discussion

Comparison of single nucleosides revealed that purines (G and A) have more pronounced affinities for the fluid lipid bilayer than pyrimidines (C and U). However, the forces driving the association are disparate. G adsorbed parallel to the membrane surface where it participated in multivalent hydrogen bonding thanks to its several hydrogen bond donating groups. Adenosine binding, conversely, was powered not by formation of energetically favourable interactions with the membrane, but by sequestration of the nucleobase from the water molecules into the membrane layer (the so called hydrophobic effect). Furthermore, we observed that the binding affinity with the membrane increased with the strand length only for polyG. For other polynucleotide sequences, including polyA, we could not observe statistically significant variation in *P* s or Δ*F* s as a function of the sequence length. Finally, we observed that unfolded RNA strands, where nucleobases have more freedom to interact with the membrane, have a higher affinity for the membrane as compared to folded RNA structures of the same length, where instead nucleobases are involved in base pairing interactions.

Although our simulations provide important takeaways for comprehending RNA-lipid interactions, certain inherent limitations need to be considered. First, the applied force fields, *χ*OL3 [26] and Lipid21 [27], have been parameterised to adequately describe RNA and membrane systems separately, but have not been extensively benchmarked for mixed systems containing both RNA and lipid molecules. Thus, it is not clear how faithfully they are able to describe RNA-membrane interactions. However, comparison to available experimental data suggests the selected force fields are able to capture the core features of RNA-membrane association. Most importantly, experimental studies corroborate preferential binding of G-rich RNA sequences to the membrane [12]. Additionally, investigations into adsorption of nucleic acids to the DPPC monolayer in the absence of divalent metal ions demonstrated that single-stranded RNA strongly associates with fluid zwitterionic lipid monolayers, while no significant adsorption of double-stranded RNA was observed, in line with our results [16]. Second, our simulations were performed without divalent metal ions. These may affect RNA binding modes and affinities, since their presence may unlock a new type of RNA-membrane interaction where phosphate groups from the RNA backbone and polar lipid head-groups jointly coordinate a divalent metal ion. Remarkably, while experiments identified G as the primary interacting nucleotide also in presence of Mg^2+^ ions, base pairing had a positive effect on the binding efficiency, opposite to that observed in systems lacking divalent ions [12]. Similarly, RNA-membrane association depends also on the composition of the lipid bilayer. Indeed, it was shown that inclusion of cationic lipids increases RNA binding [19]. As such, parsing out the effect of membrane composition and divalent ions on RNA-membrane association via MD simulations is a compelling avenue for future research. However, accurately simulating divalent metal ion ions poses a challenge as standard force fields do not account for the dynamic redistribution of divalent cations’ electronic charge that occurs when they interact with water molecules or ligands. This leads to inaccurate descriptions of key properties, such as the strength and geometry of cations’ hydration shells and their ligand-binding properties [34]. Additionally, the timescales for the exchanges in magnesium hydration shell are of the order of microseconds, making sampling challenging.

In spite of these limitations, our simulations elucidate the molecular basis driving RNA-membrane association. We conclude that in the absence of positively charged environment, owing to divalent metal ions and charged lipids, the RNA-membrane interaction is ushered by hydrogen bonding between nucleobases and zwitterionic lipid head groups and further identify the guanine nucleobase as the principal actor in RNA membrane interactions. Accordingly, we show that organisation of RNA molecules driven by base pairing decisively reduces the membrane binding affinity. Taken together, the outcomes of our simulations suggest the presence of disordered G-rich segments may be a hallmark of membrane binding RNA molecules. Our work has important implications for understanding RNA-membrane association in living organisms, for the design of RNA-based drugs and lipid-based RNA delivery systems.

## Methods

### Simulation parameters

Simulations were performed with *χ*OL3 force field [24–26] for RNA molecules, Lipid21 [27] for DPPC membranes, Joung-Cheatham parameters [35] for monovalent ions, along with the TIP3P water model [36]. Hamilton’s equations of motion were integrated using a 2 fs time step. Electrostatic interactions were evaluated using the particle mesh Ewald method [37], with an electrostatic cutoff of 1 nm. Van der Waals cutoff was set at 1 nm. Temperature was fixed to 323 K using the velocity rescaling thermostat with the coupling time constant of 1 ps [38]; pressure was set at 1 bar using a semi-isotropic cell-rescaling barostat [39] with the coupling time constant of 5 ps. The surface tension parameter (*γ*) was tuned so as to enforce the experimental average area per lipid. Formally, this can be obtained by applying the maximum entropy principle [40] to enforce the average area per lipid while minimizing the changes in the generated ensemble. By trial and error, we determined that a surface tension *γ* =100 bar·nm translates to an area per lipid of around 63.5 Å^2^, which is in line with experimental results [41, 42]. Having a correct area per lipid is crucial for properly modeling interactions at the membrane surface. Molecular dynamics simulations were performed using GROMACS 2022.3 [43, 44]; PLUMED 2.8.1 [45, 46] was used for metadynamics and trajectory analysis purposes. Additional simulation details are in Table S1.

#### Metadynamics simulations of nucleoside and polynucleotides

We generated membrane bilayers via CHARMM-GUI [47, 48]. The obtained membrane was relaxed for 100 ns for the smaller patch and for 300 ns for the bigger systems by performing unbiased molecular dynamics simulations in the NP*γ*T ensemble. Number of water molecules and lipids for the different systems are written in Table S1. Next, RNA molecules were placed 3.5 nm away from the center of the DPPC membrane along the z-axis. Water molecules and ions overlapping with the added RNA molecule were removed. We neutralised the systems with Na^+^ counter-ions and added NaCl at 150 mM concentration. All RNA/membrane systems were equilibrated for at least 100 ns by performing unbiased molecular dynamics. Next, to simulate the binding/unbinding of RNA molecules to the lipid bilayer we ran metadynamics using the following collective variable (CV). At each timestep, we computed the position of the center of the nucleotide and the center of the membrane, defined as the average position of the carbon atoms located at the end of DPPC lipid tail inside the membrane. This was used to obtain the z-component of the vectorial distance *d_z_* between the center of the nucleoside or polynucleotide and the center of the membrane. This distance was then divided by the height of the box *L_z_* in that timestep, and multiplied by the value of the average height of the box *< L_z_ >*, to obtain the effective z-distance *d̃*_*z*_ that served as the CV for biasing. This was done to enforce periodicity of the bias between the two sides of the membrane. The fluctuations of the height of the box were very small (less than 2%), therefore the error involved in this practice is negligible. During analysis and in all the reported plots we used the real z-distance *d_z_*, and not the effective *d̃*_*z*_. Simulation lengths and metadynamics parameters are listed in Table S1.

### Metadynamics simulations of a 19-mer RNA strand

We simulated the sequence *rAGGGAAGGGAAGGGAAGGG* whose membrane binding properties were previously examined experimentally [12]. The sequence is assumed to form a G-quadruplex in the presence of K^+^ ions, however no structural data is available. Therefore, the G-quadruplex structure was obtained by homology modelling. As template, we used the 5’ untranslated region of neuroblastoma RAS gene (PDBID: 7SXP) [49] to model the guanosine core of the G-quadruplex, containing two K^+^ ions, while completely removing the loops and the first two nucleotides. We manually added the three *AA* loops by using the two-nucleotide loop topology from the X-ray structure of the B-raf domer (PDBID: 4H29) [50]. Next, we solvated the structure, performed energy minimisation with restraints on the guanine core and collected unbiased molecular dynamics in plain water for almost 2 *µ*s. Sugar puckering at the final configuration inserted on the membrane showed prevalence of C3’-*endo* on the loops, even if the initial structure of the loops was from a DNA structure. The equilibrated G-quadruplex structure was then used to perform metadynamics simulations. The geometrical center of the G-quadruplex was inserted 4.5 nm away from the center of the membrane. We used two CVs for biasing. The first CV was *d̃*_*z*_, defined as above. To calculate the G-quadruplex center, only the central guanines were used. The second CV was the cosine of the angle (cos(*θ*)) between the central axis of the G-quadruplex and the normal to the membrane (parallel to the z-axis of the simulation box). We used cos(*θ*) rather than the *θ* itself because identically independently distributed unitary vectors lying on a sphere display a uniform distribution in cos(*θ*). To be certain that the G-quadruplex did not unfold during the metadynamics simulation where strong biasing forces are applied, we added a harmonic wall to the distance between the two K^+^ located in the guanine core. To unbias for the effect of these restraints, this contribution was used when reweighting the frames. For simulations of the 19-mer in the unfolded state, the initial structure was built using Avogadro [51]. The unfolded RNA strand was then solvated and simulated in water for almost 1.5 *µ*s. A representative structure, shown as inset in Fig. 4b, was manually chosen among those obtained by performing a cluster analysis and used in subsequent metadynamics simulations. The RNA/membrane system was then initialised similarly as in the case of the G-quadruplex structure. At each timestep, we calculated for each phosphorus atom of the RNA strand, its minimum distance from all the phosphorus atoms of the membrane. The first biased CV was the minimum of these distances, while the second CV was the maximum value among the minimum distances. This allowed the strand to both attach and detach from the membrane. For analysis purposes, we used the minimum and maximum distances from the membrane center, defined as above.

### Partition coefficient calculation

After simulating the systems, we defined the functions *f ^L^* and *f ^W^* for each *i_th_* frame. These equal 1 when the RNA molecule is in the membrane bound or unbound state, respectively. The bound and unbound states were defined by four z-distance values *d*_1_*, d*_2_*, d*_3_ and *d*_4_. The parameters relevant for the definition of the bound state were *d*_1_ and *d*_2_. They were chosen in such a way that *F* (*d_z_*) had its minimum value for *d*_1_ *< d*^∗^ *< d*_2_. The parameters which defined the unbound state were *d*_3_ and *d*_4_, which were chosen such that *F* (*d_z_*) ≈ const for *d*_3_ *< d_z_ < d*_4_. We then computed the partition coefficient *P* as:

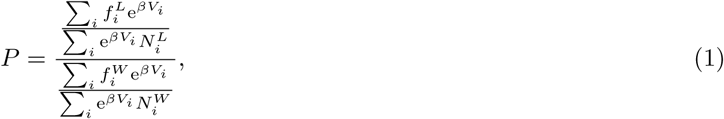

where *V_i_*was the value of the bias in the *i_th_*frame of the simulation and *β* = <u>^1^</u> . We simplified the expression by noting that the number of lipids present in the simulation (*N^L^*) was constant. The number of water molecules (*N^W^*) was instead approximated by using the density of TIP3P water (*ρ*_wat_ ≈ 0.997 <u>^kg^</u>). We then simplified the expression as follows:

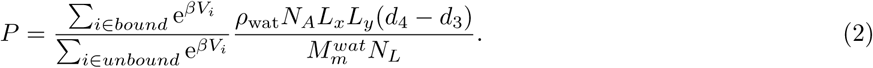

In the case of the RNA strand, we used *d*_min_ instead of |*d_z_*| to define the bound and unbound state.

### Error estimation

Errors were estimated in two separate ways. We exploited the fact that the free energy profile at convergence should be symmetric *F* (*d_z_*) = *F* (−*d_z_*). We thus obtained two independent estimates for all observables, occurring at two separate sides of the membrane. The first contribution to the error was obtained by computing the deviation between these two independent estimates. The second, smaller, source of error was estimated by performing Bayesian Bootstrap1. [52] separately for each side of the membrane. The two sources of error were then combined as independent error sources.

### Contact analysis

Hydrogen bonds were computed via MDAnalysis [53, 54], using 3.5 Å as a distance threshold between the hydrogen bond donor and acceptor heavy atoms and 130° for the angle formed between the hydrogen bond donor, proton and acceptor. Van der Waals contacts analysis was also performed by using MDAnalysis. Two heavy atoms were assumed to form a van der Waals contact if their interatomic distance was less than the sum of the two van der Waals radii plus 0.6 Å.

## Acknowledgments

We thank CINECA for the provided computational hours which allowed us to perform this work. We thank Tomás F.D. Silva and Pavlína Pokorná for helpful discussions and feedback. A.M. and J.A. thank the PNRR: National Center for Gene Therapy and Drugs based on RNA Technology CUP B83C22002860006 CN_0000004.

## Supplementary information

**Table S1:**
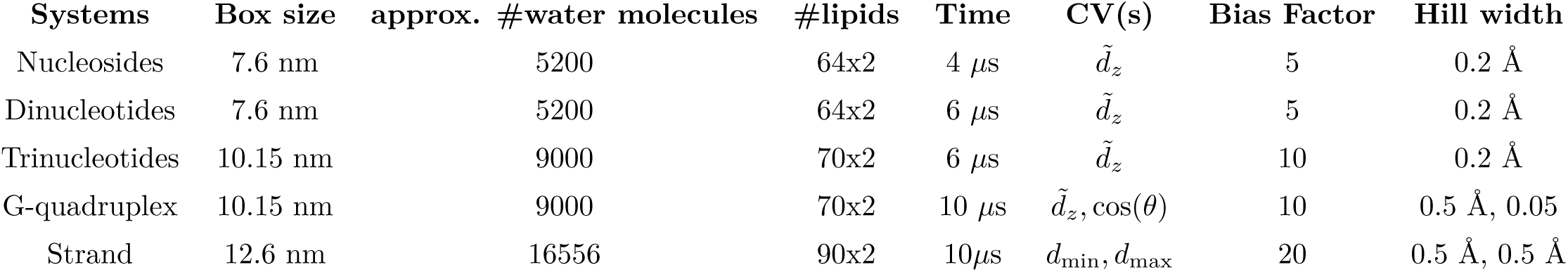
Relevant parameters used for the different systems. *d̃*_*z*_ indicates the effective z-distance from the membrane center (see Methods).

| **Systems** | **Box size** | **approx. #water molecules** | **#lipids** | **Time** | **CV(s)** | **Bias Factor** | **Hill width** |
| --- | --- | --- | --- | --- | --- | --- | --- |
| Nucleosides | 7.6 nm | 5200 | 64x2 | 4 μs | $\tilde{d}_z$ | 5 | 0.2 Å |
| Dinucleotides | 7.6 nm | 5200 | 64x2 | 6 μs | $\tilde{d}_z$ | 5 | 0.2 Å |
| Trinucleotides | 10.15 nm | 9000 | 70x2 | 6 μs | $\tilde{d}_z$ | 10 | 0.2 Å |
| G-quadruplex | 10.15 nm | 9000 | 70x2 | 10 μs | $\tilde{d}_z, \cos(\theta)$ | 10 | 0.5 Å, 0.05 |
| Strand | 12.6 nm | 16556 | 90x2 | 10 μs | $d_{\min}, d_{\max}$ | 20 | 0.5 Å, 0.5 Å |

**Figure S1:**
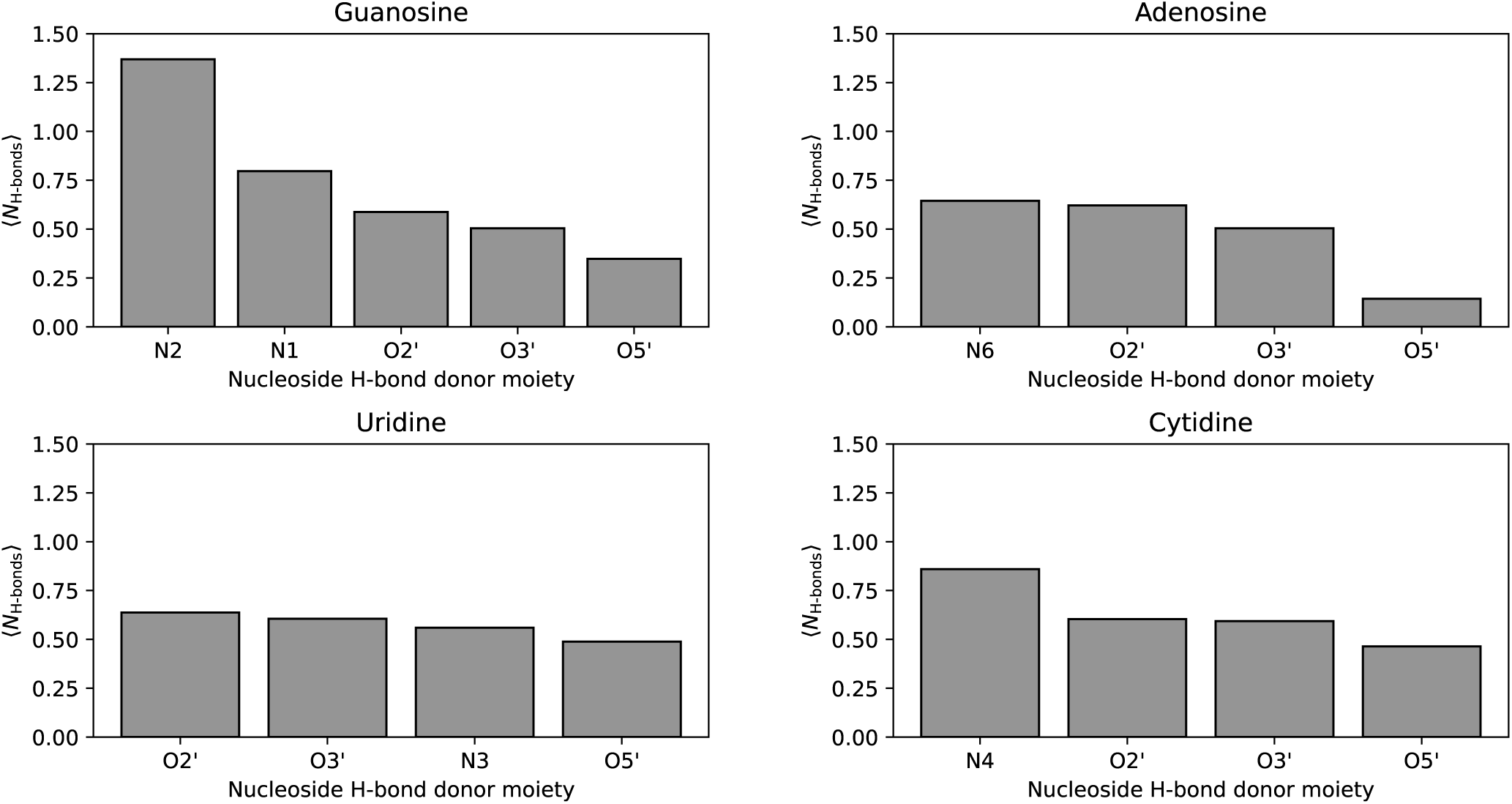
Average hydrogen bonding frequencies ⟨*N*_H-bonds_⟩ for each hydrogen bond donating moiety in nucleosides. DPPC lipids do not have hydrogen bond donating moieties, thus only RNA donors are shown.

**Figure S2:**
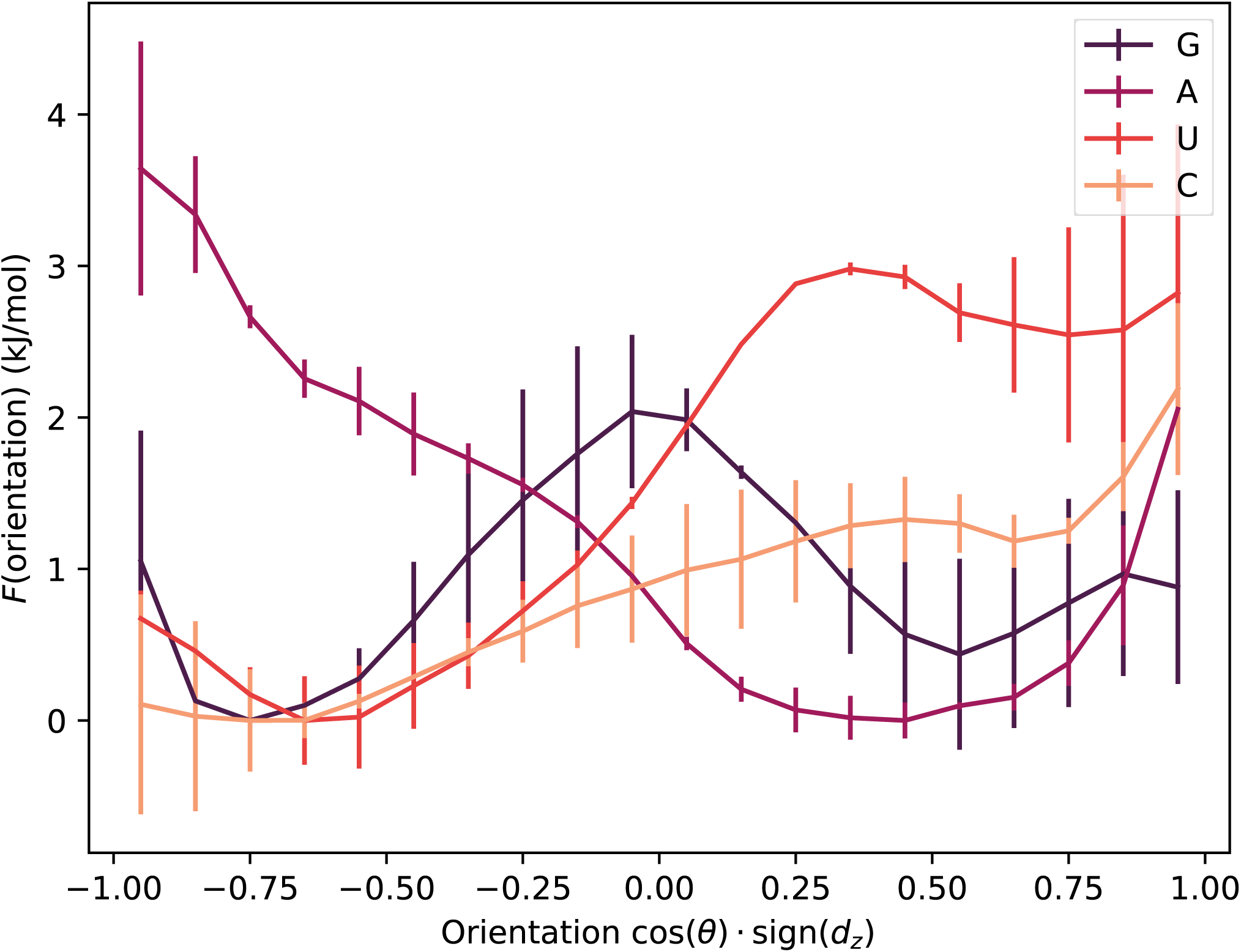
Free energy *F* as a function of the orientation of the nucleobase (cos(*θ*) · sign(*d_z_*)) on the membrane for the different nucleosides. Multiplying cos(*θ*) by sign(*d_z_*) is done to achieve equivalence between the upper and lower membrane leaflet. G tends to be parallel to the membrane surface, whereas A is usually perpendicularly inserted inside the membrane. The absence of symmetry between positive and negative cosines is due to the presence of the ribose and syn/anti asymmetries of different nucleobases.

**Figure S3:**
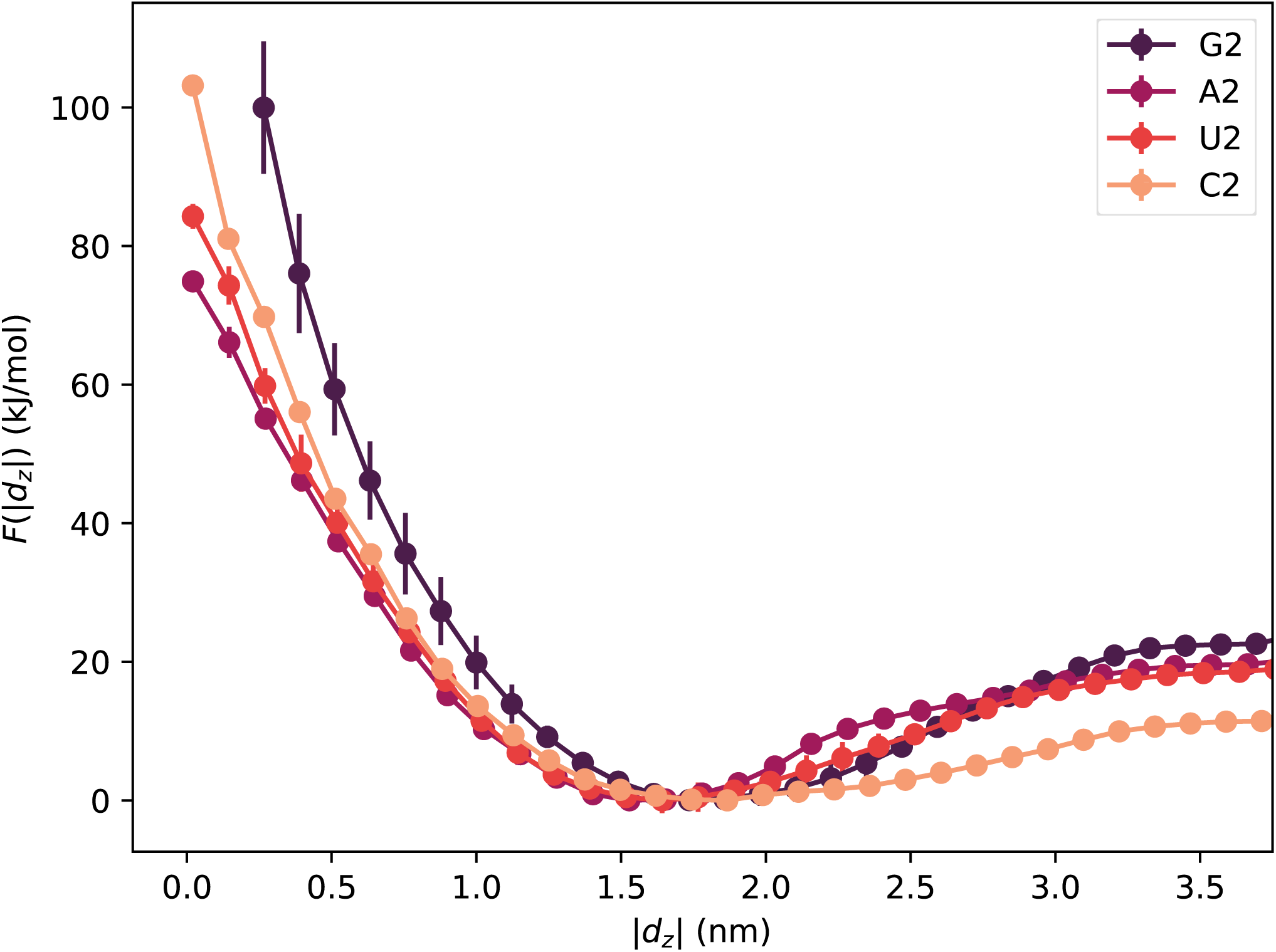
Free energy *F* as a function of the absolute value of the z-distance between the membrane center and the geometric center of dinucleotides |*d_z_*|.

**Figure S4:**
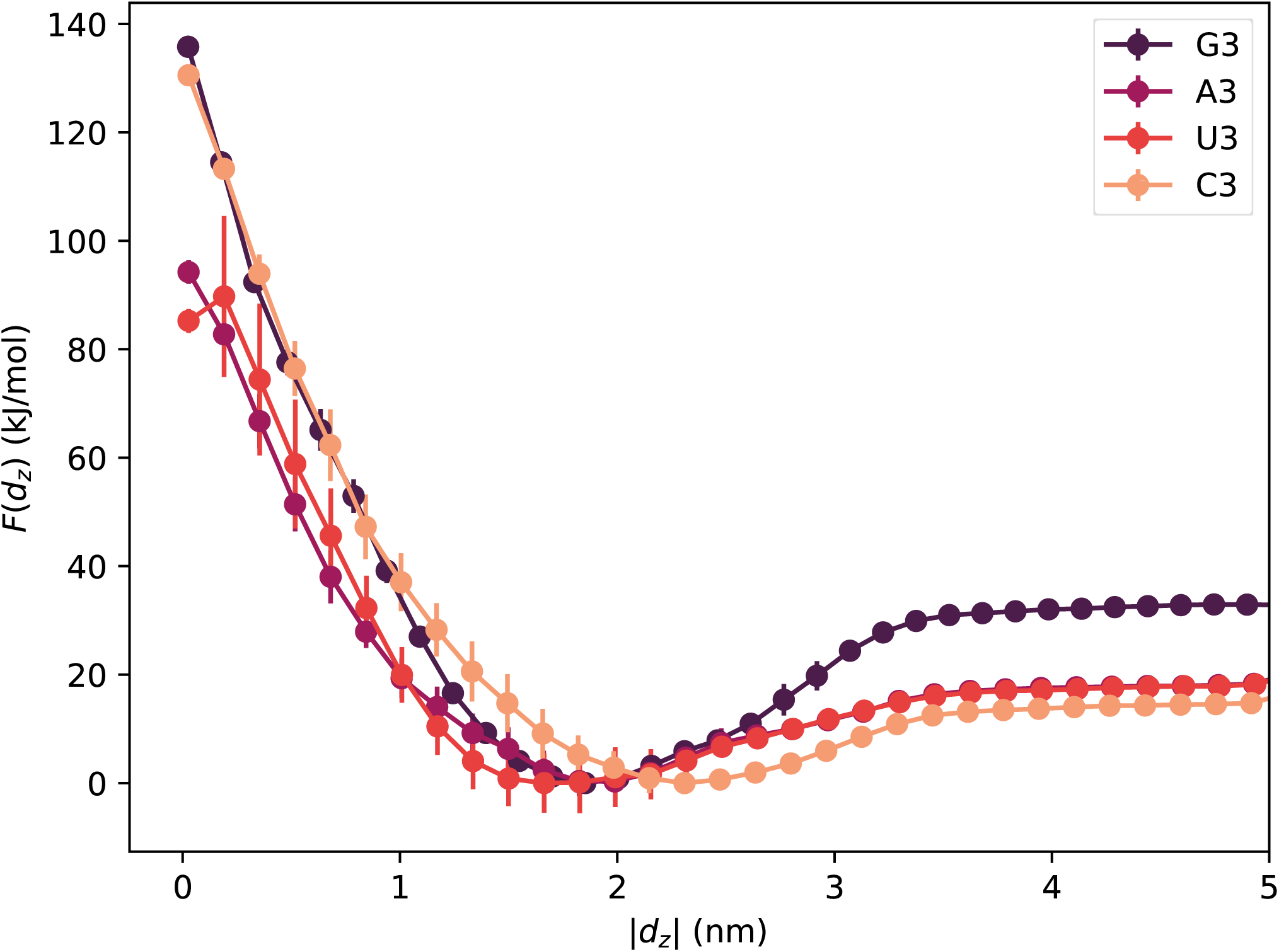
Free energy *F* as a function of the absolute value of the z-distance between the membrane center and the geometric center of trinucleotides |*d_z_*|.

**Figure S5:**
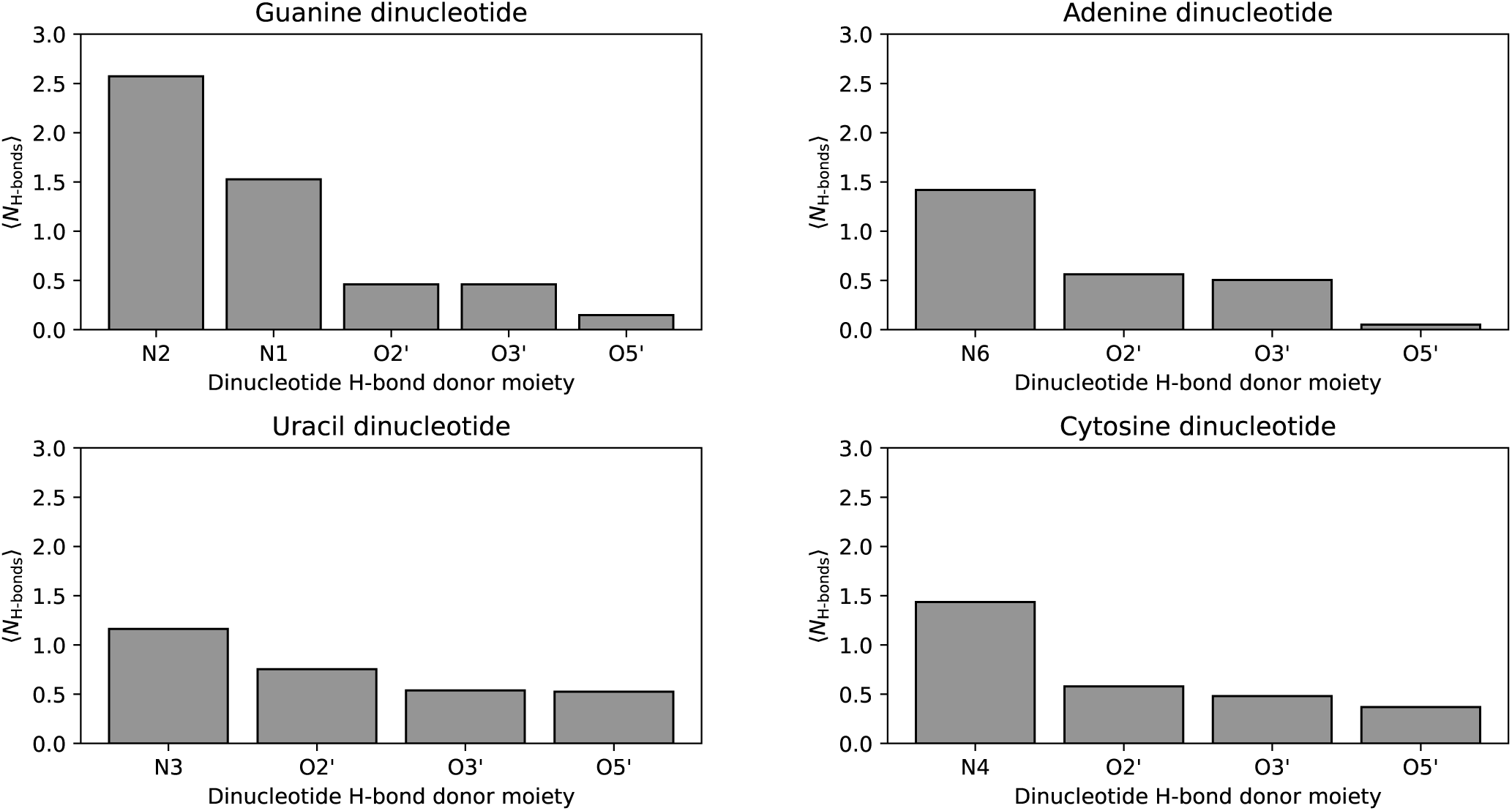
Average hydrogen bonding frequencies ⟨*N*_H-bonds_⟩ for each hydrogen bond donating moiety in dinucleotides.

**Figure S6:**
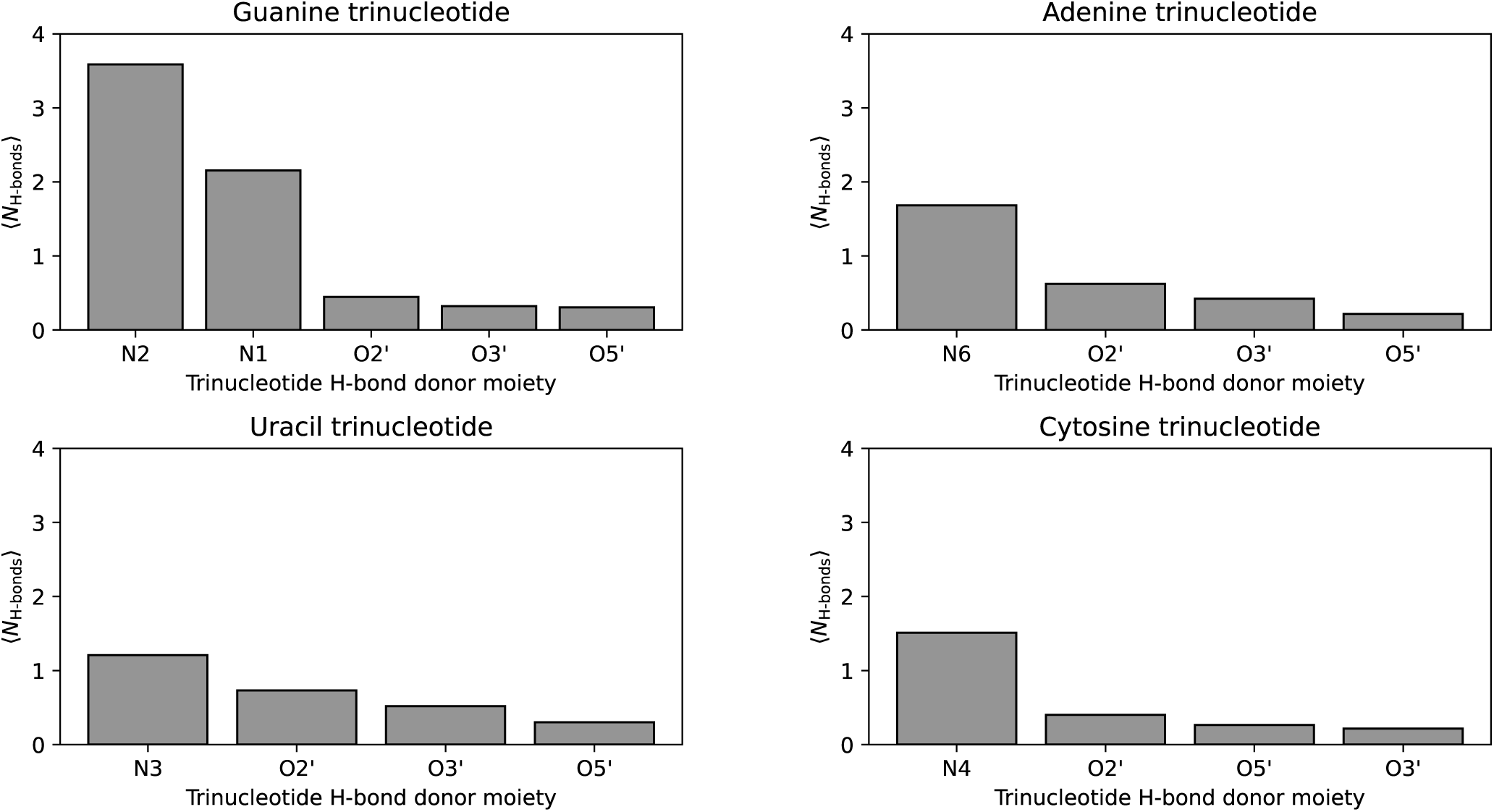
Average hydrogen bonding frequencies ⟨*N*_H-bonds_⟩ made by each hydrogen bond donating moiety in trinucleotides.

**Figure S7:**
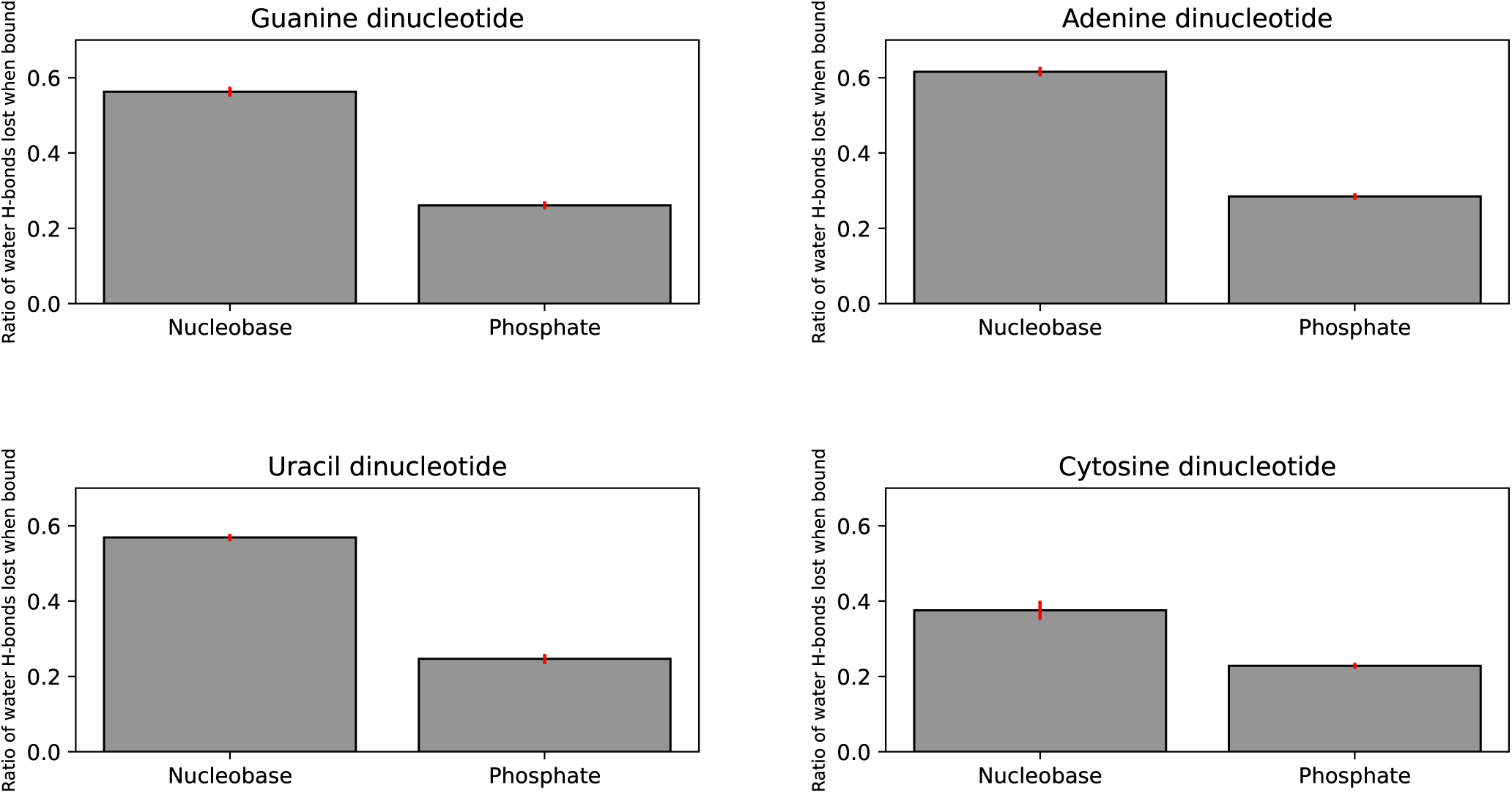
Percentage of water hydrogen bonds lost upon dinucleotide binding to the membrane for the nucleobases and phosphate groups. The percentage was obtained by calculating the difference between the number of hydrogen bonds the dinucleotides form with water in the bound and unbound state and dividing it by the average number of water hydrogen bonds made in the unbound state. It can be seen that phosphate moieties seem to retain most of their hydrogen bonds with water molecules, while more than 50% of water hydrogen bonds with nucleobases are lost upon binding, with the exception of the cytosine dinucleotide, which loses almost 40%.

**Figure S8:**
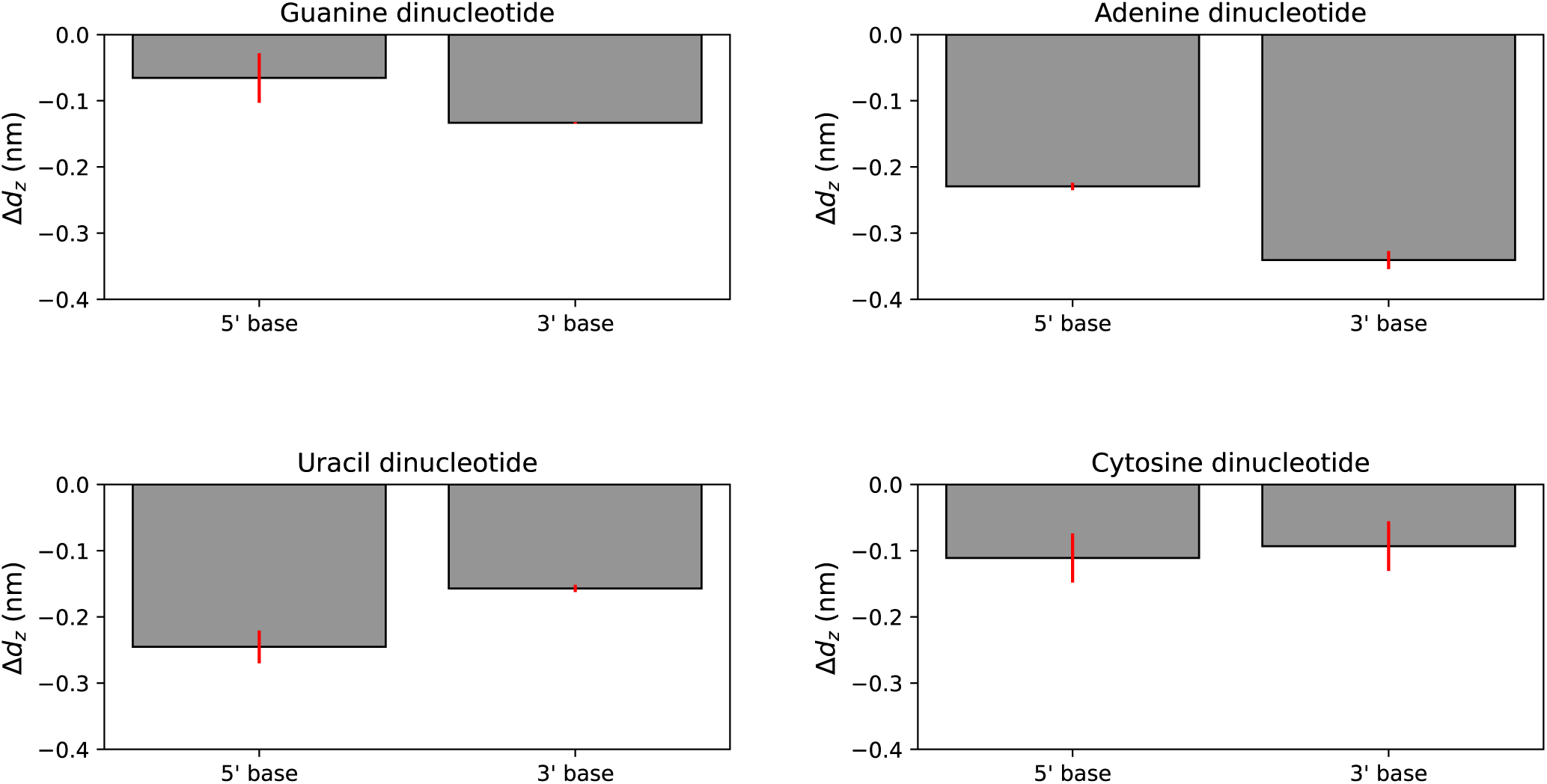
Difference between |*d_z_*| of the center of mass of 5’ and 3’ nucleobases and the phosphorus atom in the phosphate group (Δ*d_z_* = |*d_z,_*_base_| − |*d_z,P_* |) for the dinucleotides in the bound state. It can be seen that the phosphate is on average positioned further away from the membrane surface than the two nucleobases and prefers to interact with the solvent.

**Figure S9:**
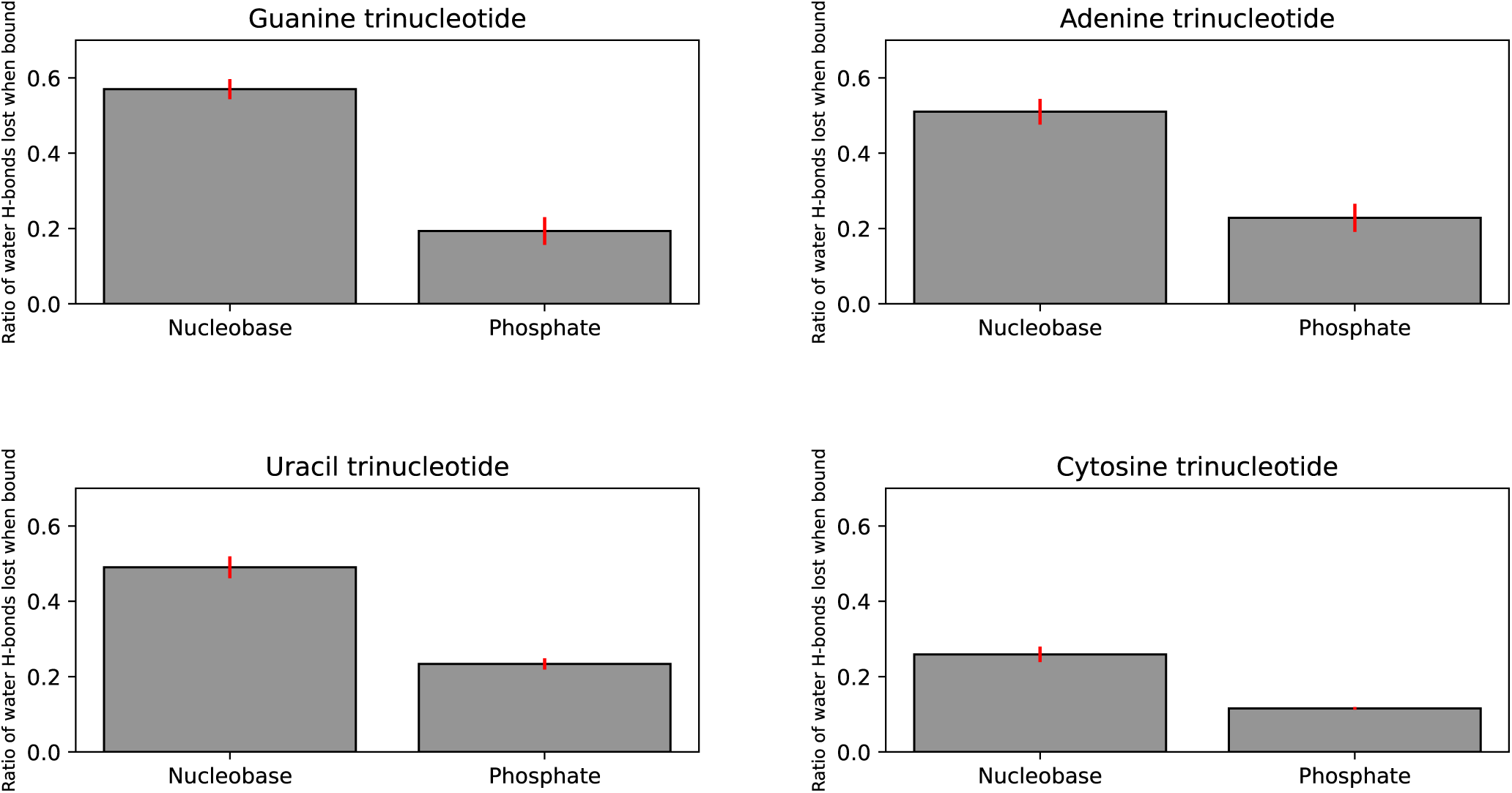
Percentage of water hydrogen bonds lost upon trinucleotide binding to the membrane for the nucleobases and phosphate groups. In comparison to nucleobases, the phosphate moieties retain significantly more hydrogen bonds with water.

**Figure S10:**
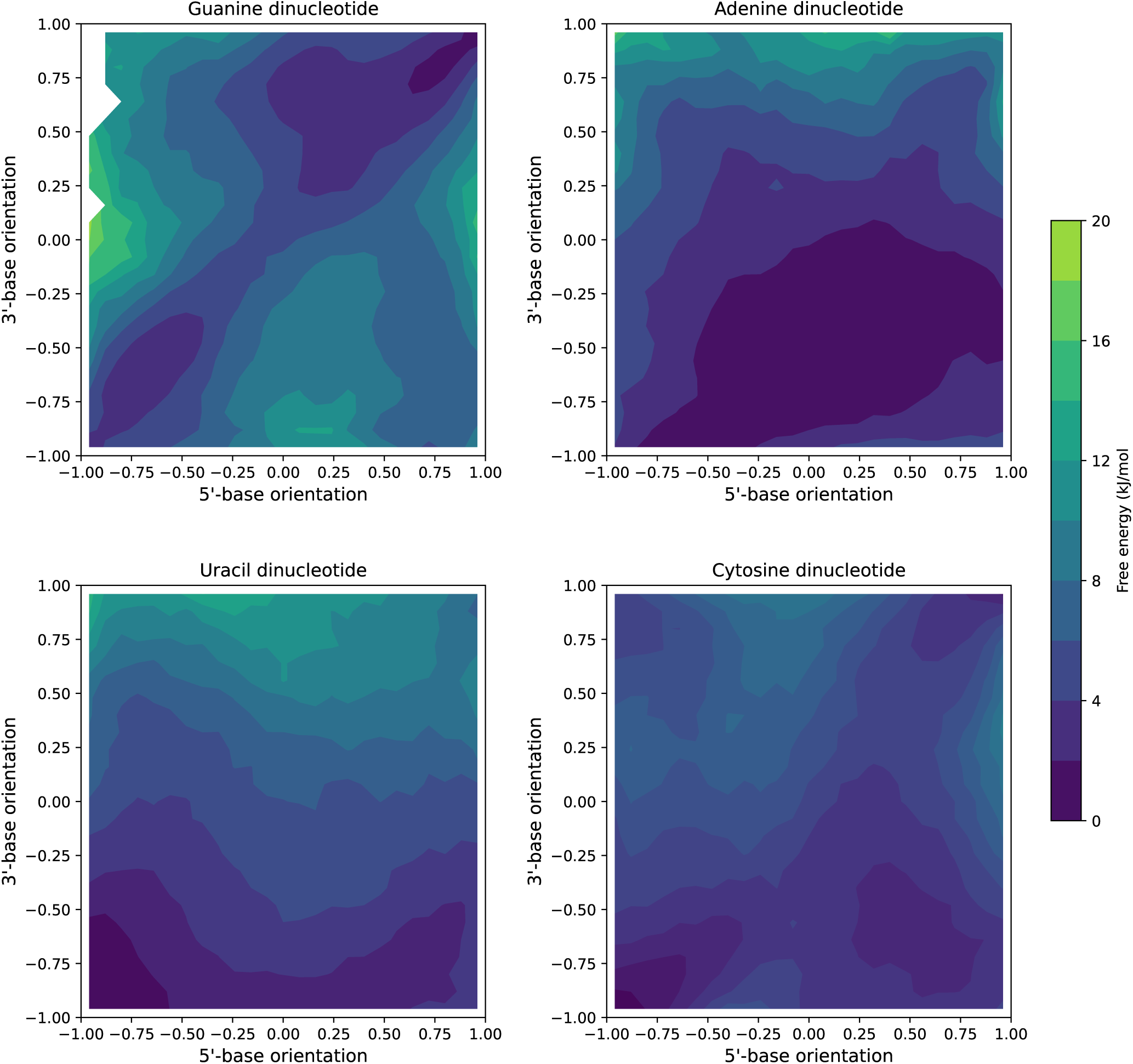
Nucleobase orientations (cos(*θ*) · sign(*d_z_*)) for both nucleobases with respect to the membrane normal for dinucleotides. In case of guanine, uracil and cytosine dinucleotide, both nucleobases preferred to orient parallel to the membrane. Conversely, in case of the adenine dinucleotide there was a broader range of optimal orientations. One nucleobase was preferentially inserted, while the other assumed diverse orientations with respect to the membrane.

**Figure S11:**
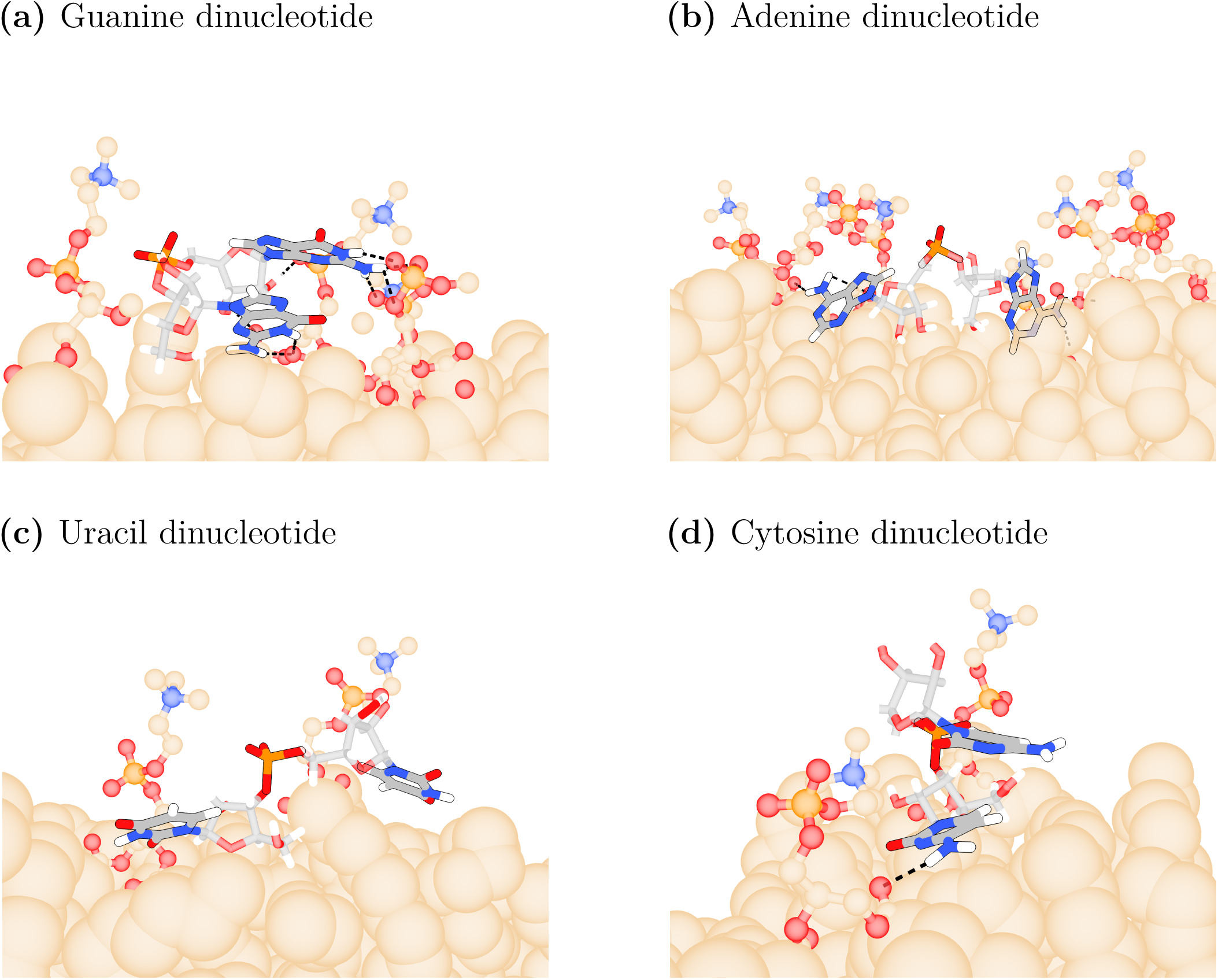
Snapshots of dinucleotides in free energy minimum showing likely orientations with respect to the membrane. Only lipid head groups forming hydrogen bonds or van der Waals contacts to nucleotides are displayed. Hydrogen bonds are displayed with dashed black lines.

**Figure S12:**
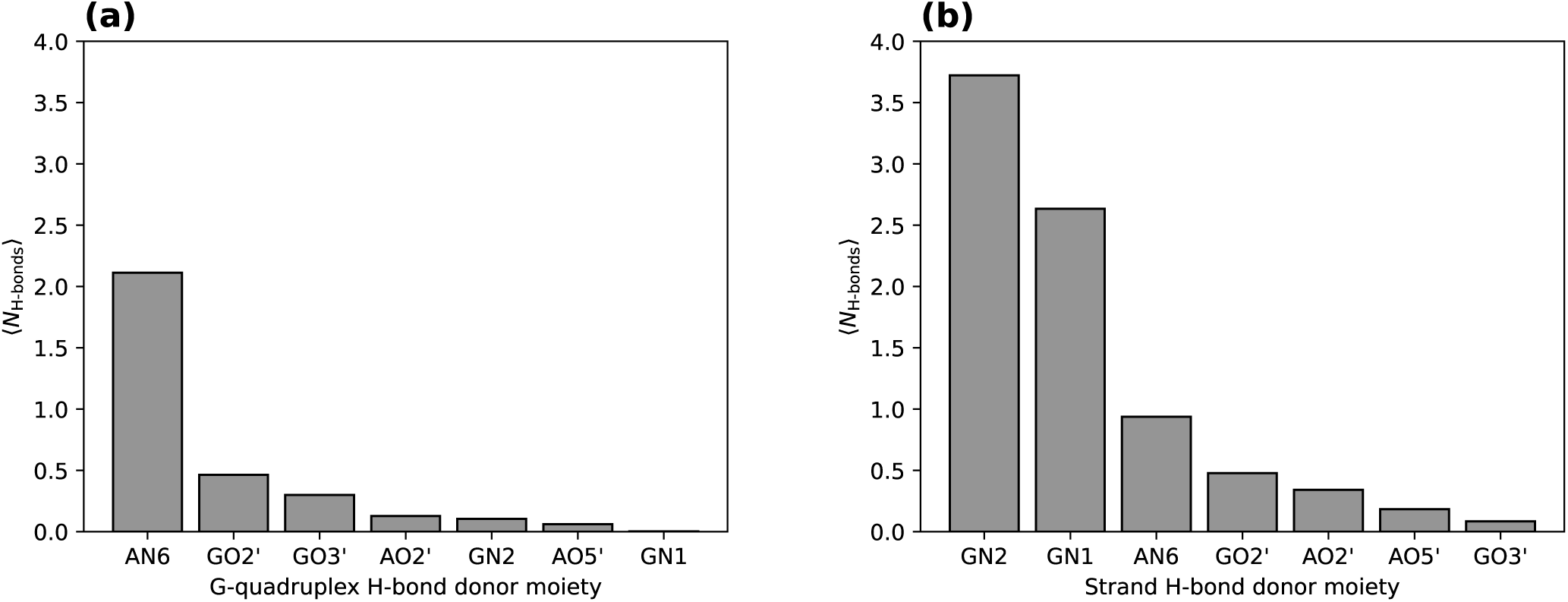
Average number of hydrogen bonds (⟨*N*_H-bonds_⟩) for each hydrogen donor moiety, for the G-quadruplex

**Figure S13:**
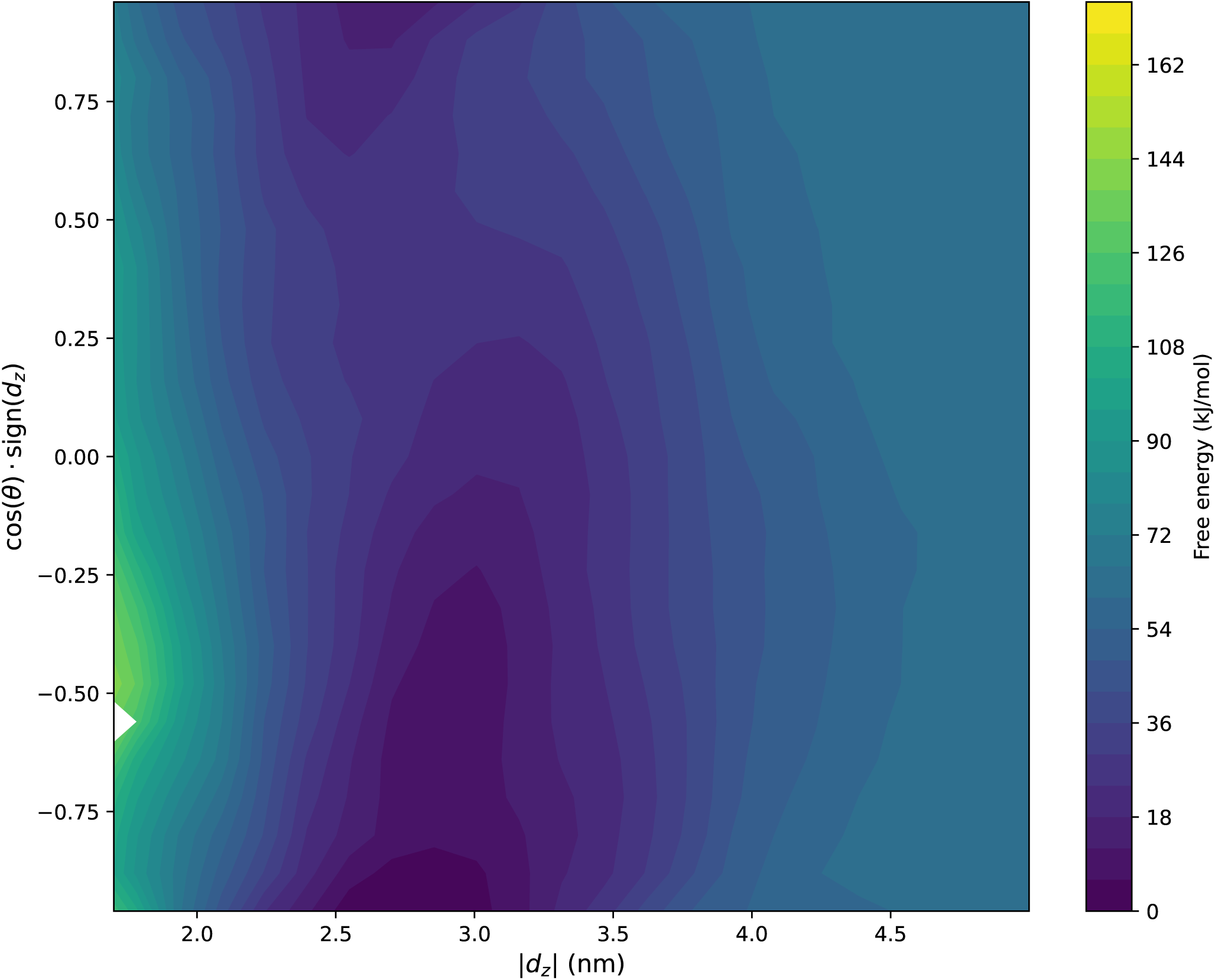
G-quadruplex free energy, as a function of its orientation with respect to the membrane cos(*θ*) · sign(*d_z_*) and the absolute value of the z-distance |*d_z_*|.

**Figure S14:**
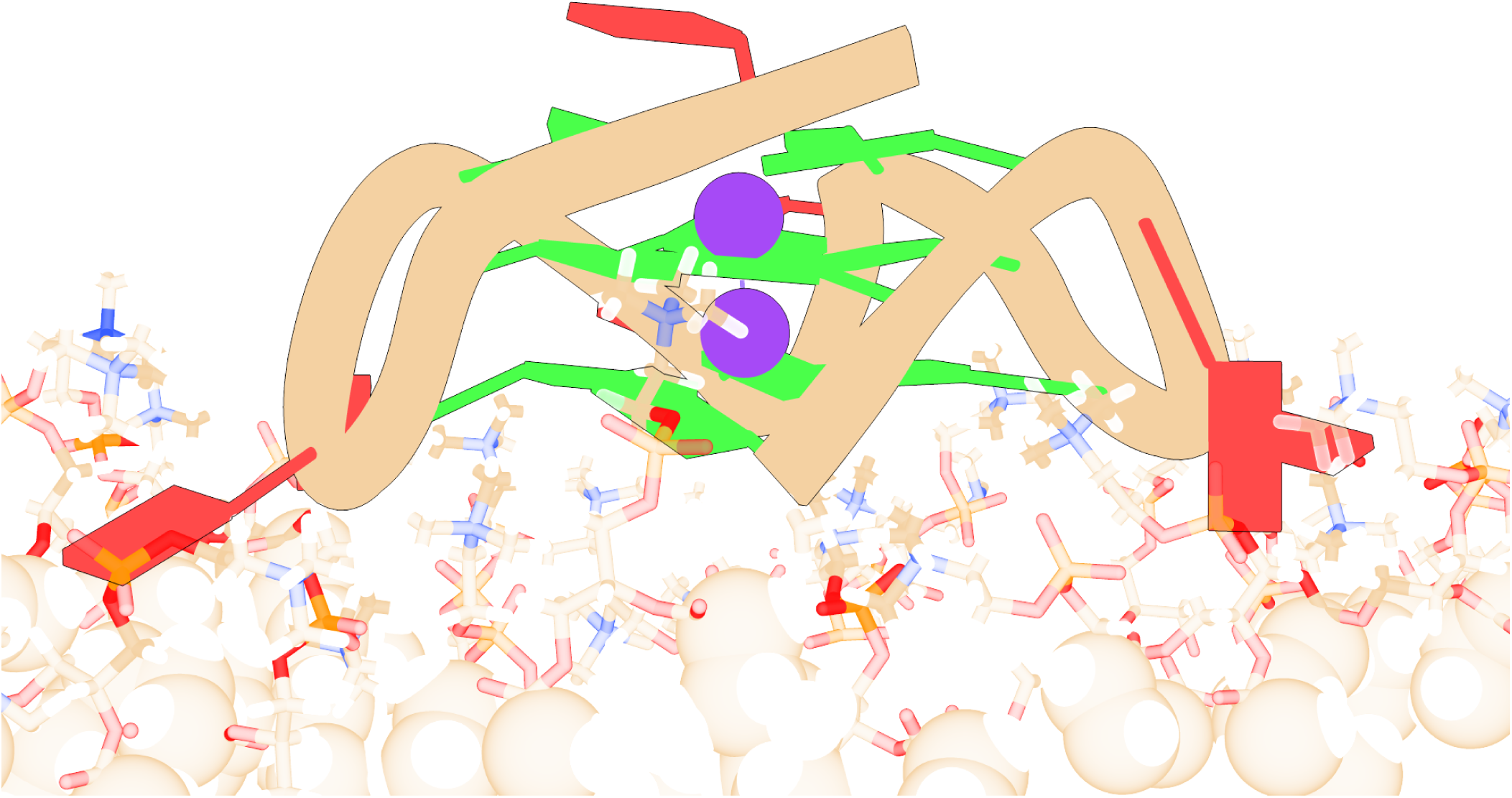
Membrane-bound state of G-quadruplex in its most likely orientation. Guanines are represented in green, adenines in red.

**Figure S15:**
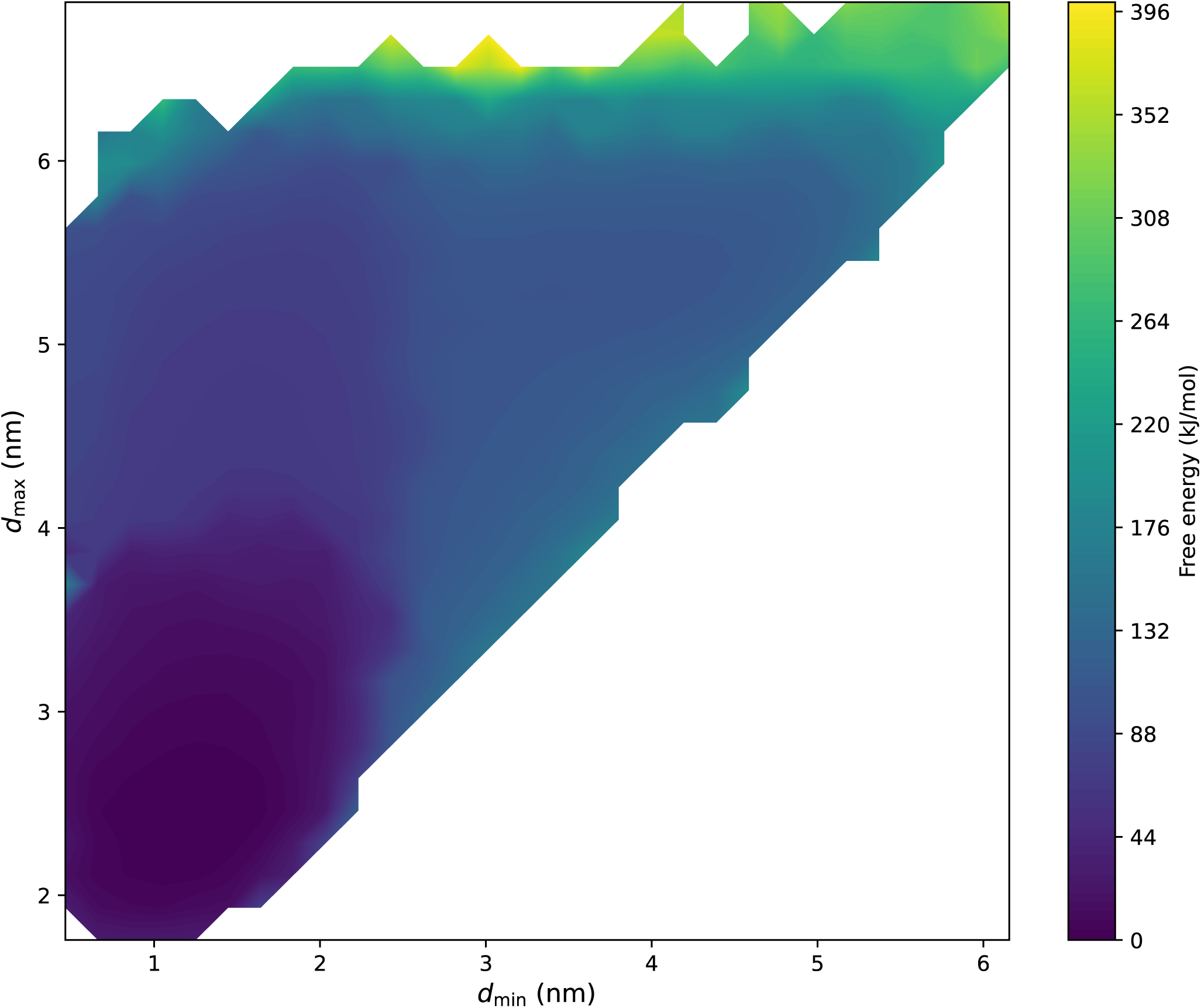
Free energy of the unfolded RNA strand as a function of the minimum (*d*_min_) and maximum (*d*_max_) distance between the RNA strand phosphates and the membrane center (see Methods).

**Figure S16:**
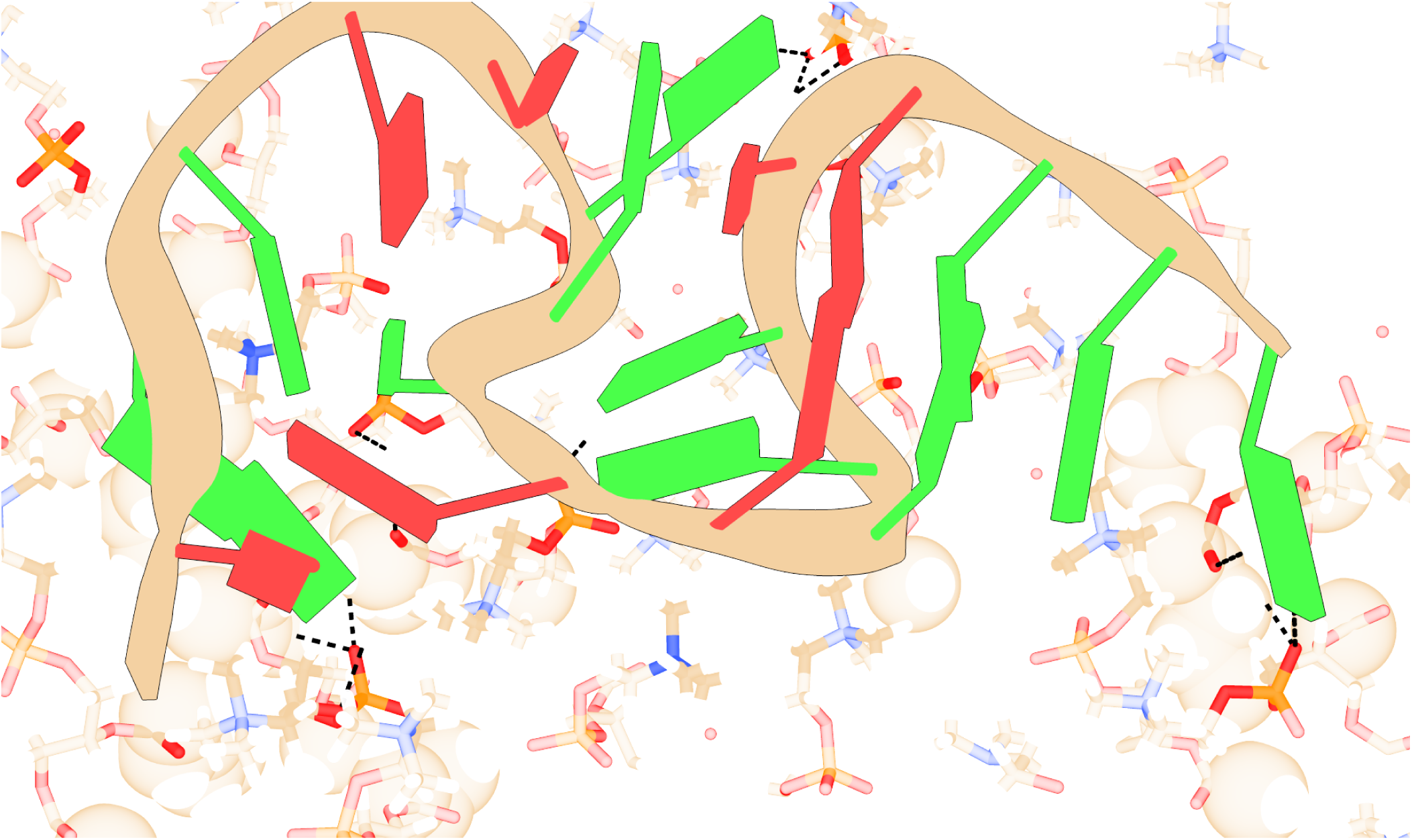
Top-view of the unfolded RNA strand bound to the membrane, extracted from the free energy minimum. Guanines are shown in green and adenines in red. Interaction with the membrane is primarily mediated by hydrogen bonding (black dashed lines) to guanine residues.

## Notes

### Competing Interest Statement

The authors have declared no competing interest.

